# ReMeDy: A Flexible Statistical Framework For Region-based Detection of DNA Methylation Dysregulation

**DOI:** 10.64898/2026.01.02.697394

**Authors:** Suvo Chatterjee, Siddhant Meshram, Ganesan Arunkumar, Fasil Tekola-Ayele, Arindam Fadikar

## Abstract

Region-based epigenome-wide association studies have demonstrated improved statistical power and biological interpretability compared with probe-wise analyses of DNA methylation data. However, most existing region-based methods characterize methylation dysregulation primarily through changes in mean methylation levels associated with a phenotype of interest. Substantial evidence indicates that phenotype-associated methylation alterations may also manifest through changes in methylation variability or through joint shifts in mean and variability. Despite this, no existing statistical framework jointly models mean–variance methylation changes in a region-based manner. We propose ReMeDy, a flexible statistical framework that uses a hierarchical likelihood approach within a generalized linear model setting to identify differentially methylated regions, variably methylated regions, and regions exhibiting joint differential and variable methylation at a genome-wide scale. Unlike existing models, ReMeDy operates directly on biologically defined co-methylated regions, allowing it to naturally capture spatial correlation inherent in DNA methylation array data, while avoiding reliance on heuristic, user-defined tuning parameters such as smoothing spans and kernel bandwidths that can substantially influence results and introduce subjectivity. Through extensive simulation studies and comprehensive benchmarking against popular models, we demonstrate that ReMeDy maintains false discovery and type-I error rates at nominal levels while achieving consistently higher statistical power across a wide range of realistic scenarios. Application to population-level DNA methylation data further shows that ReMeDy identifies biologically meaningful regions and pathways implicated in complex human diseases that are not captured by conventional mean-based analyses alone. ReMeDy is implemented as an open-source R package and is freely available at https://github.com/SChatLab/ReMeDy.

## 1 Introduction

DNA methylation (DNAm) is a central epigenetic mechanism involved in the regulation of gene expression, chromatin organization, and cellular identity, serving as a dynamic interface between genetic variation, environmental exposures, and disease susceptibility ^1–3^. Over the past decade, array-based technologies, particularly the Illumina HumanMethylation450 and MethylationEPIC platforms, have enabled epigenome-wide association studies (EWAS) at unprecedented scale, facilitating systematic, genome-wide investigations of DNAm patterns across diverse populations and disease contexts ^4–6^. These studies have shown that disease-associated methylation changes are often not restricted to individual Cytosine-phosphate-Guanine (CpG) sites but instead occur across clusters of spatially correlated CpGs spanning regulatory regions such as promoters, enhancers, and CpG islands ^7–13^. Consequently, region-based analyses of methylation data have emerged as a powerful alternative to single-site testing, offering increased statistical power, a reduced multiple-testing burden, and improved robustness to technical noise ^11;14;15^. Genomic regions identified using this region-based approach are referred to as differentially methylated regions (DMRs), defined as regions in which the mean methylation levels of CpGs differ significantly between groups or treatment conditions. Importantly, substantial evidence indicates that DMRs tend to exhibit greater biological interpretability and improved reproducibility across cohorts, making them more suitable for downstream functional and translational analyses and for identifying potential therapeutic biomarkers ^16–18^.

Despite substantial advances in study design, preprocessing, and statistical methodology, a major conceptual bias persists in EWAS, where epigenetic dysregulation is still predominantly characterized by differences in mean DNAm between phenotypic groups or exposure categories ^18–20^. Most widely used EWAS pipelines are explicitly optimized to detect shifts in average methylation levels at individual CpG sites or aggregated genomic regions ^21;22^, implicitly assuming that within-group variability is homogeneous or incidental to the primary scientific question, and rarely treating variance effects as explicit targets of inference ^23–25^. Although mean-based frameworks have proven effective for identifying robust and reproducible associations, they impose inherent limitations on the classes of epigenetic dysregulation that can be detected ^26^. Recent studies highlight that methylation dysregulation is more appropriately conceptualized as comprising three distinct components, mean, variance, and joint mean–variance changes, and that disease-associated alterations may manifest through not only mean methylation changes but also through increased dispersion, heterogeneity, or epigenetic instability even in the absence of substantial mean shifts ^27–29^. Such variability-driven effects are increasingly recognized in complex and heterogeneous conditions, including cancer, autoimmune diseases, and aging-related phenotypes ^28;30–34^. Consistent with recent studies, we provide additional corroborative evidence in this study demonstrating that all three components represent integral components of the methylation landscape (Figure 1), and therefore must be jointly investigated to avoid an incomplete and potentially biased understanding of epigenetic mechanisms underlying disease susceptibility and progression ^23;26^.

**Figure 1:**
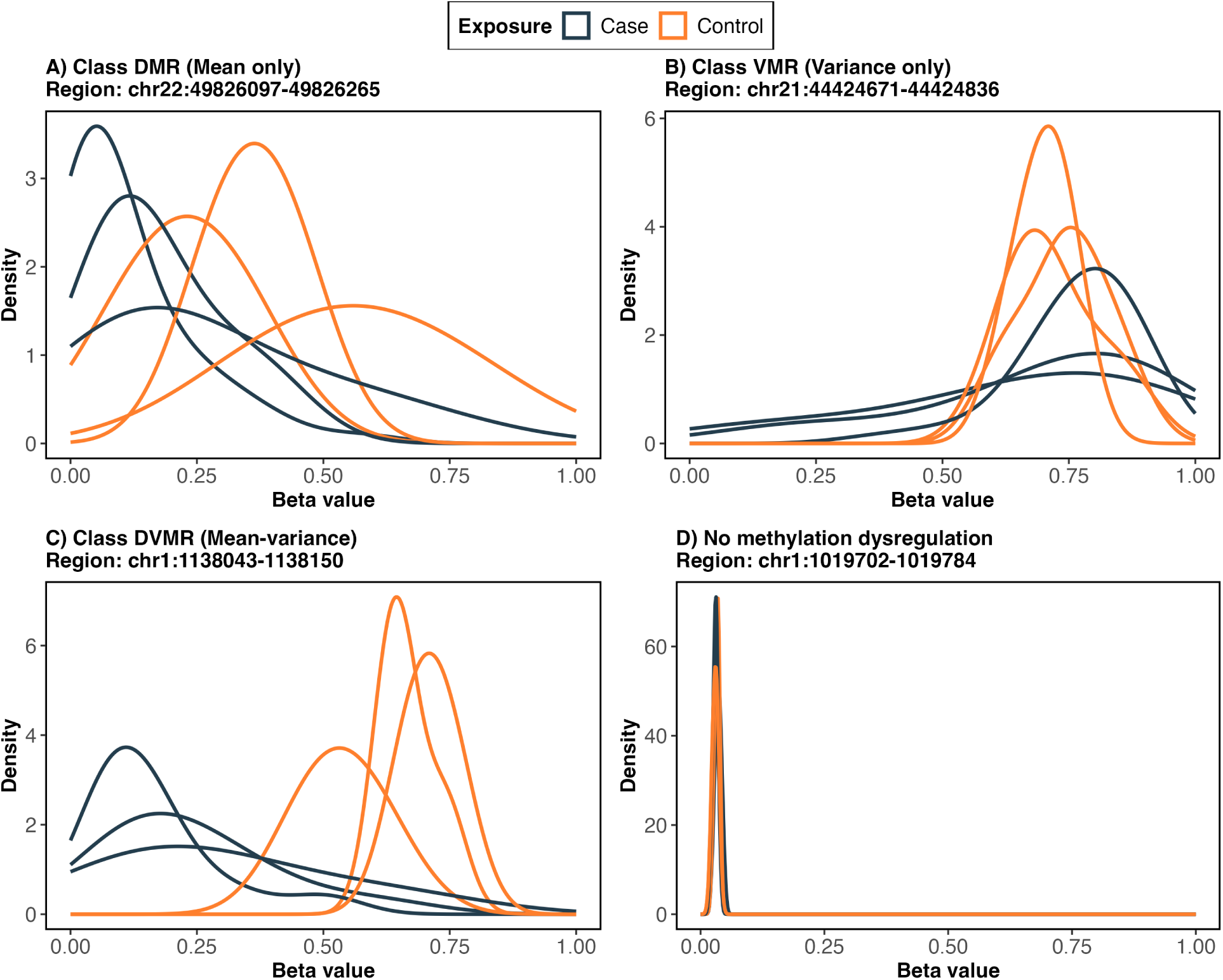
**This figure illustrates distinct components of methylation dysregulation observed across the genome**. Randomly selected regions are shown to demonstrate four patterns of methylation change between cases and controls, visualized using beta value density distributions. In each panel, the three density curves per group correspond to the three CpG sites within the region. Panel A shows a DMR, characterized by a location shift, where the density peaks are displaced between groups, indicating differences in mean methylation levels. Panel B shows a VMR, where the mean methylation levels are similar between groups but the spread of the distributions differs, reflecting changes in methylation variability. Panel C shows a DVMR, where both a location shift and changes in variability are observed. Panel D shows a region with no methylation dysregulation, where the distributions overlap between cases and controls.

Existing methods for detecting methylation dysregulation, including both differential methylation (DM) and variable methylation (VM) can be best viewed as a collection of methodological design choices rather than cohesive inferential frameworks, resulting in a fragmented analytical landscape ^35–37^. Region-level DM approaches typically rely on outcome-guided region discovery, aggregating probe-wise association statistics across neighboring CpGs using smoothing, clustering, or p-value combination methods ^15;38;14^. While these strategies leverage local spatial correlation to improve power, they are sensitive to tuning parameters, prone to outcome-driven boundary overfitting, and often identify regions that do not reflect intrinsic epigenomic organization, thereby reducing generalizability and interpretability ^14;39;40^. In addition, these summary-statistic based approaches inherits the assumptions, biases, and calibration properties of underlying probe-wise tests and depends on accurate estimation of spatial autocorrelation, which can be unstable in modest sample sizes or under irregular probe spacing common to methylation data ^39;41^. Methodological limitations are even more pronounced for VM analysis. Firstly, existing VM models are mostly site-based approaches and ignore the spatial dependence structure inherent to DNAm data ^23;42^. Secondly, variance is typically modeled independently of the mean, despite fundamental mean–variance coupling in methylation data which can lead to mis-specified variance estimates and fragmented biological interpretation ^43^. Thirdly, VM methods have restricted modeling flexibility, as most frameworks do not accommodate covariates to adjust for confounding or technical artifacts (such as batch effects, cell-composition shifts) and often limit the primary exposure to binary designs, substantially limiting their applicability to more complex epidemiological studies ^23;44;45;42^. Collectively, this highlights that methylation dysregulation continues to be characterized in a piecemeal fashion, and that the absence of integrated region-level frameworks capable of jointly modeling location and scale effects warrants the development of a unified statistical framework that can capture all components of methylation dysregulation while accounting for the structural and biological complexity of DNAm data.

To address these methodological gaps, we propose ReMeDy (integrated statistical framework for <u>Re</u>gion-based detection of <u>Me</u>thylation Dysregulation), which identifies differentially methylated, variably methylated, and joint differential–variable regions within a single coherent statistical framework. Firstly, ReMeDy is built on a hierarchical likelihood approach within a generalized linear model setting, enabling region-level modeling of methylation data which accounts for correlation among neighboring CpG sites through random effects and providing principled inference under spatial dependence. Secondly, unlike existing region-based methods that rely on outcome-guided region aggregation and are therefore sensitive to tuning parameters and boundary overfitting, ReMeDy operates only on co-methylation regions defined independently of the phenotype, ensuring that regions reflect intrinsic epigenomic organization and improving interpretability and generalizability. Thirdly, the framework performs joint mean–variance modeling, explicitly accounting for the mean–variance dependence inherent to methylation data, and thereby avoids the mis-specified variance estimates and biased inference that can arise when location and scale effects are modeled separately. Lastly, ReMeDy supports covariate/confounder adjustments, incorporation of multiple random effects to capture technical and biological sources of variation, and accommodates binary, continuous, and discrete phenotypic variables. Collectively, these features position ReMeDy as an integrated and statistically principled framework that addresses the fragmented nature of existing approaches and enables comprehensive characterization of methylation dysregulation in array-based EWAS. To our knowledge, this represents the first systematic implementation of a hierarchical generalized linear modeling (HGLM) framework with joint mean–variance modeling for identifying phenotype-associated methylation markers in DNAm data.

The remainder of the manuscript is organized as follows. In Section 2, we first describe the hierarchical likelihood framework within the generalized linear model (GLM) setting and introduce the resulting HGLM in the context of DNAm array data, along with algorithms for estimating model coefficients under a joint mean–variance modeling framework. In Section 3, using both synthetic data and published methylation array datasets, we conduct a systematic benchmarking study of more than ten representative methods for detecting aberrant methylation. We show that our proposed method, ReMeDy, consistently outperforms existing approaches across experimental platforms in terms of statistical power and false discovery rate control. Finally, Section 4 concludes with a brief discussion of our findings. Our open-source software package is available at https://github.com/SChatLab/ReMeDy.

## 2 Methods

### 2.1 Current modeling schemes of DNAm array data

DNAm array data exhibits complex dependence structures arising from both biological and technical sources. CpG sites are spatially organized along the genome and frequently display correlated methylation patterns due to shared regulatory mechanisms, chromatin organization, and genomic proximity ^7;46;47^. Several strategies have been proposed to address correlation in DNAm data. A widely used class of models perform CpG-level association testing followed by post hoc aggregation of nearby significant sites using smoothing ^38^, p-value combination and autocorrelation ^48^ based approaches. These approaches leverage spatial proximity to implicitly account for local correlation but do not model dependence directly within the statistical framework. A less commonly used class of methods applies classical models for correlated data, such as linear mixed models (LMMs) ^49^ and generalized estimating equations (GEEs) ^50^, which explicitly model spatial dependence by treating clustered CpGs as repeated measurements within individuals ^51;52;47^. While these approaches represent important progress in handling correlation in DNAm data, a common shortcoming still persists. Despite growing evidence that methylation dysregulation can manifest through changes in mean, variance, or their joint behavior, most existing methods are primarily formulated to assess mean effects and provide limited support for joint mean–variance modeling ^27–29^. This serves as our motivating factor to develop a statistical framework that can jointly capture correlation in DNAm data along with mean-variance changes in methylation patterns at the regional level.

### 2.2 Hierarchical likelihood approach for modeling correlated methylation data

Likelihood-based inference forms the foundation of many statistical models, but classical likelihood approaches encounter challenges when applied to correlated and hierarchical data. In LMMs, random effects are introduced to account for correlation, and under Gaussian assumptions the random effects can be integrated out analytically to yield a closed-form marginal likelihood ^49^. However, they are primarily designed to model mean structure, with variance components treated as nuisance parameters rather than quantities directly linked to covariates ^49^. As a result, LMMs offer limited flexibility for modeling systematic changes in variability across phenotypic groups. Generalized linear mixed models (GLMMs) ^53^ extend mixed-effects modeling to non-Gaussian responses, but likelihood evaluation requires integration over random effects, which is analytically intractable and typically relies on numerical approximations such as Laplace methods or quadrature ^54^. While these approximations are often adequate for modeling mean effects, they become less reliable and increasingly computationally demanding when variance parameters are allowed to depend on covariates, limiting their practical use for joint mean–variance modeling ^54^. GEEs work with a marginal likelihood approach by specifying a working correlation structure, but they do not define a full likelihood, limiting inference on variance components and likelihood-based model comparison ^50^. To overcome these limitations, Lee and Nelder introduced the hierarchical likelihood (h-likelihood) approach ^55;56^. Rather than integrating out random effects, the h-likelihood defines a joint likelihood for the observed data and the unobserved random effects, treating the random effects as additional unknown quantities to be estimated alongside the model parameters. This formulation explicitly represents correlation through shared random effects while retaining a coherent likelihood-based framework. By avoiding high-dimensional integration, the h-likelihood enables simultaneous estimation of fixed effects, random effects and dispersion parameters, making it suitable for modeling correlated biological data such as DNAm.

### 2.3 Hierarchical generalized linear modeling scheme for DNAm data

The h-likelihood framework leads to HGLMs which extends GLMMs by allowing both the mean and dispersion components of the response distribution to be modeled through separate but linked linear predictors, each of which may include fixed effects and random effects ^57^. This structure enables joint inference on mean and variability while explicitly accounting for correlation. Random effects are introduced at the subject level to account for correlation among CpGs measured within the same individual, treating CpG sites within a region as repeated measurements. At the same time, dispersion model allow variability to depend on biological outcomes, enabling direct modeling of variable methylation effects. Together, this framework not only accounts for the correlation in the measurements but also allows methylation differences linked to a phenotype of interest to be characterized through changes in mean methylation, methylation variance, or both, within an integrated statistical framework.

### 2.4 Proposed Model Specification for Region-Based DNAm Analysis

In this study, we propose to model region-level DNAm data using the h-likelihood approach under the HGLM framework. Let 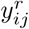 denote the methylation level for sample *i* (*i* = 1*, . . . , N*) at CpG site *j* (*j* = 1*, . . . , n_r_*) within genomic region *r* (*r* = 1*, . . . , R*). Since methylation levels are analyzed on the logit-transformed scale (M-values), which approximately follow a Gaussian distribution, we assume a Gaussian likelihood with an identity link, i.e.,

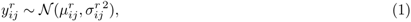

where 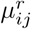 denotes the conditional mean of the methylation level for CpG site *j* in sample *i* modeled through the mean model, and 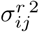 denotes the residual variance, which is modeled through the variance model described in the following sections. These two models are linked within the HGLM framework. The mean model characterizes changes in mean methylation levels while the variance model captures changes in methylation variability that is linked to the phenotypic variable of interest which can be binary, continuous or discrete in nature. Together, these components enable joint inference on mean and variance within an integrated h-likelihood formulation.

#### 2.4.1 Mean Model

For a given genomic region *r*, the conditional mean of the methylation level 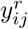 at CpG site *j* for sample *i*, given the subject-specific random effect *b_i_*, is modeled as

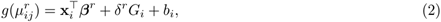

where *g*(·) denotes the identity link and 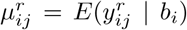. The covariate vector **x***_i_* includes an intercept and any additional adjustment variables, with corresponding region-specific coefficients ***β****^r^*. The parameter *δ^r^* represents the region-specific phenotype-associated effect on mean methylation, with *G_i_* denoting the phenotype variable of interest. Statistical inference on *δ^r^* therefore provides a direct region-level assessment of DM, and regions with significant *δ^r^* are identified as DMRs. The subject-specific random intercept 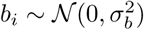 accounts for correlation among CpG sites measured within the same individual, treating CpG sites within region *r* as repeated measurements.

#### 2.4.2 Variance Model

To model methylation variability across samples and phenotype of interest, the residual variance 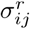 is modeled using a GLM with a log link,

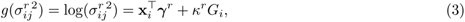

where 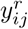 denotes the residual variance of the methylation level at CpG site *j* for sample *i* within region *r*. The covariate vector **x***_i_* includes an intercept and any additional adjustment variables, with corresponding region-specific coefficients ***γ****^r^*. The parameter *κ^r^* represents the phenotype-associated effect on methylation variability for region *r*, with *G_i_* denoting the phenotype of interest. Statistical inference on *κ^r^*therefore provides a direct region-level assessment of VM and regions with significant *κ^r^*are identified as VMRs.

This two-part mean–variance formulation allows for simultaneous modeling of DM (mean differences) and VM (variance differences) across genomic regions. In this study, although our focus has been on a binary (case-control) phenotype variable, however, this proposed framework can trivially accommodate continuous and discrete phenotypic variable of interest.

### 2.5 Estimation via h-likelihood

Estimation of the model defined in equations (1)-(3) requires inference on the region-specific fixed effects ***β****^r^* and ***γ****^r^*, the phenotype of interest effects *δ^r^* and *κ^r^*, the subject-level variance component 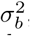, and the subject-specific random effects 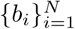 . Let 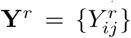 denote the observed CpG-level M-values within genomic region *r*. Typically, classical likelihood-based inference is based on the marginal likelihood obtained by integrating out the random effects,

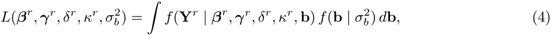

where **b** = (*b*_1_*, . . . , b_N_*)^⊤^. Given the high-dimensional nature of DNAm data, direct evaluation of the marginal likelihood in (4) can be computationally unstable and demanding due to the presence of random effects and the explicit modeling of variance.

The h-likelihood framework ^55;57^ avoids explicit integration by treating the random effects as additional unknown quantities to be estimated jointly with the fixed effects. The h-likelihood for region *r* is defined as

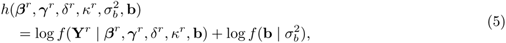

which combines the conditional likelihood of the data given the random effects with the log-density of the random effects implied by their assumed Gaussian distribution. For the Gaussian HGLM considered here, the conditional log-likelihood contribution from observation (*i, j*) is

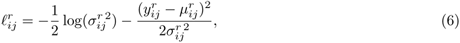

where 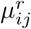 and 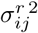 are defined by the mean and variance models in equations (2) and (3), respectively.

Maximization of the h-likelihood proceeds through an iterative procedure that alternates between updating the mean and variance components. Conditional on current values of the variance parameters, maximization of (6) with respect to the mean structure leads to weighted estimating equations equivalent to fitting a linear mixed model with weights 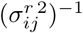. This step yields updated estimates of ***β****^r^*, *δ^r^*, and the subject-specific random effects **b**. Conditional on the updated mean structure, information about the variance parameters is contained in the squared residuals 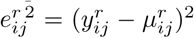. Writing 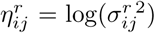, differentiation of (6) with respect to 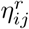 shows that the score for the variance model depends on the quantity 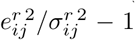, which motivates treating the squared, standardized residuals from the mean model as the response for updating the variance model. These mean and variance updates are alternated until convergence.

### 2.6 Hypothesis testing and multiple testing adjustment

Inference for DM and VM is conducted at the region level using Wald-type test statistics derived from the fitted HGLM. For each genomic region *r*, let 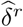 and 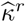 denote the estimated region-specific phenotypic effects in the mean and variance model, respectively. Let 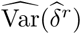 and 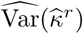 denote the corresponding variance estimates obtained from the h-likelihood–based estimation procedure. DM is assessed by testing the null hypothesis 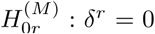 using the Wald statistic 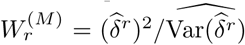, which is asymptotically distributed as 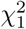 using 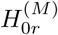. The resulting *p*-value is denoted by 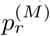. Similarly, VM is assessed by testing 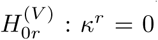. using the Wald statistic 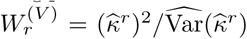, which is also asymptotically 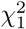, yielding a variability-related *p*-value 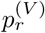. Here the superscript ‘M’ denotes mean model and ‘V’ denotes variance model. To assess joint evidence for regions exhibiting both DM and VM, we combine the two marginal *p*-values using the Cauchy Combination Test (CCT) ^58^. For each region *r*, we define the test statistic

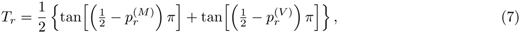

and compute the corresponding joint *p*-value as

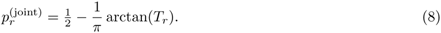

The CCT yields valid combined *p*-values under arbitrary dependence between 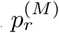 and 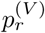.

To account for multiple testing across genomic regions, the sets 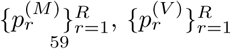 and 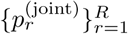 are adjusted separately using the Benjamini–Hochberg (BH) procedure ^59^ to control the false discovery rate (FDR). Regions with adjusted 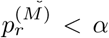 (*α* denotes a pre-specified statistical significance threshold such as *α* = 0.05.) are declared DMRs, regions with adjusted 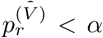 are declared VMRs, and regions with adjusted 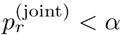 are declared differentially and variably methylated regions (DVMRs).

## 3 Results

First, we describe the practical implementation details of ReMeDy, including model formulation and input requirements (Section 3.1). Second, we provide details about our simulation framework and the different simulation scenarios considered in our simulation framework (Section 3.2). Third, we present extensive simulation studies designed to evaluate the performance of ReMeDy under realistic methylation array data settings, comparing its ability to detect DMRs, VMRs, and DVMRs against existing methods (Section 3.3). Finally, we apply ReMeDy to analyze a publicly available DNAm array dataset to demonstrate its practical utility and to uncover biologically relevant methylation dysregulation that are not readily captured by existing approaches (Section 3.4).

### 3.1 Implementation Details

ReMeDy operates on user-supplied co-methylated regions and assumes that DNAm beta-values within these regions have been appropriately preprocessed using standard, well-established pipelines. Specifically, in our workflow, raw intensity data are processed using the minfi Bioconductor package ^60^, which performs sample- and probe-level quality control. This includes filtering probes with poor detection p-values, removing probes with known single-nucleotide polymorphisms or cross-reactive genomic mapping, and excluding sex-chromosome probes when appropriate. This is followed by normalization and for batch effect correction we use ComBat ^61^, an empirical Bayes method that models and adjusts for technical variation across batches, thereby removing batch-associated effects from the data. These correction for technical variation ensures that downstream analyses primarily reflect biological signal rather than technical noise. Following preprocessing, methylation levels are transformed to M-values and supplied to SACOMA ^62^, which identifies co-methylated regions. SACOMA utilizes a spatially-constrained hierarchical clustering algorithm to obtain clusters of contiguous CpG sites exhibiting correlated methylation patterns in a genome-wide manner. ReMeDy is agnostic to the specific normalization or batch-effects correction strategy used prior to co-methylation region construction, provided that major technical effects are adequately controlled. Although we use minfi for preprocessing and SACOMA for obtaining co-methylated regions, alternative preprocessing pipelines such as ChAMP ^63^ and co-methylated region construction methods such as coMethDMR ^47^ can be used.

### 3.2 Simulation Strategy and Settings

We evaluated the capability of ReMeDy in identifying DMRs, VMRs, and DVMRs in comparison to a broad variety of existing methods. For DM, comparisons were made with region-based approaches including bumphunter ^38^, comb-p ^48^, dmrff ^64^, and DMRcate ^65^, while VM performance was assessed against existing site-level methods such as JLSsc ^42^, iEVORA ^66^, DiffVar ^24^, Bartlett’s test ^67^, and Levene’s test ^68^ due to lack of region-based models. For joint differential and variable methylation, comparisons were made against JLSsc and iEVORA due to their ability to return CpGs exhibiting both effects. Although no published methods are explicitly designed to identify VMRs or DVMRs at the regional level, DMRcate was included in all three comparisons as its software documentation module mentions about its ability in detecting VMRs and DVMRs. Null (type-1 error) simulations were performed using datasets derived from two population-level methylation array datasets (See supplementary section 1.3 for details) representing low (N = 40) and high (N = 142) sample-size settings in which phenotypic labels of the samples were randomly permuted to remove any inherent associations and create new datasets with no true biological signals. Power simulations were performed using realistic synthetic DNAm datasets generated from multivariate normal distributions with compound symmetry region-specific correlation structure and mean and variance effects to represent DMRs, VMRs, and DVMRs, thereby providing known ground truth. Details of our simulation strategy and the compared models are provided in supplementary section 1.1 and 1.2 respectively. Model performance was evaluated using FDR, statistical power, Matthews correlation coefficient (MCC), and area under the receiver operating characteristic curve (AUROC) across all simulation scenarios (See supplementary section 1.5 for details). Simulation scenarios comprised of 10,000 regions with systematically varying sample sizes (N = 50, 100, and 150), mean effect sizes (0.4, 0.7, and 1.0), variance effects (with group 2 variability 1.5-, 2.5-, or 3.5-fold greater than group 1), and group proportions in 50%/50% or 30%/70% case–control splits (See STable 1 for summary). Results presented in this section focus on scenarios with a mean effect of 0.7, a variance effect of 2.5, and imbalanced design (30%/70% case-control split). Findings from all remaining simulation settings are reported in supplementary Section 2.

### 3.3 Simulation Results

Under null simulations with 10,000 regions and no true biological signals present in the simulated data, most methods exhibited reasonable type-I error control, including ReMeDy, which maintained error rates below the nominal 5% threshold for both sample sizes (Figure 2). DiffVar was the only method to show mild type-I error inflation in the smaller sample size (N = 40), while achieving appropriate error control in the larger sample size (N = 142) setting. Specifically DMRcate, comb-p, bumphunter, and iEVORA were highly conservative across both sample sizes, with median type-I error rates ranging from 0 to 0.1% while Levene’s test, ReMeDy, JLSsc, and Bartlett’s test demonstrated moderate conservatism to balanced error control, with median type-I error rates ranging from 2 to 5% across both sample size settings.

**Figure 2:**
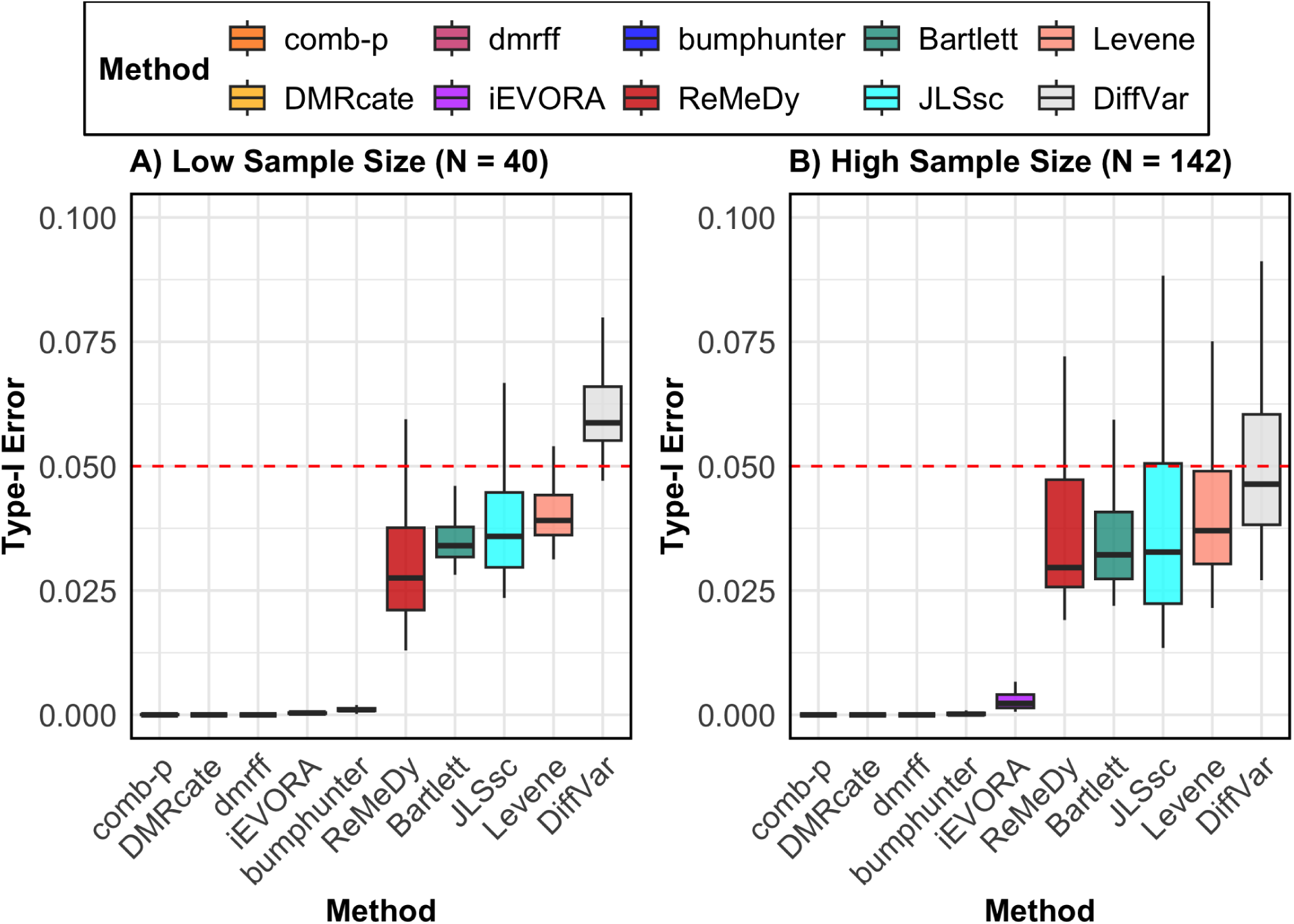
**Type-I error rate evaluation**. This panel plot illustrates the type-I error (y-axis) for ReMeDy and all competing methods under two different scenarios: Panel A shows results from a low sample size (N = 40) setting and Panel B shows results from a higher sample size (N = 142). Each boxplot represents 100 simulation runs. The dashed red line marks the nominal type-I error rate of 0.05. Methods with boxplots near or below this line are controlling false positives well. Approaches with no visible box have a type-I error rate of zero.

Beyond type-I error control, ReMeDy consistently outperformed existing DMR detection methods across all evaluation metrics under the DMR-only simulation setting, which comprised 10,000 regions, 10% true DMRs with a mean difference effect of 0.7, no variance effect, and unequal group proportions (Figure 3). ReMeDy achieved the highest statistical power across all sample sizes, with performance increasing steadily as sample size grew (Figure 3A). In comparison, DMRcate, comb-p, and bumphunter showed modest power gains, while dmrff consistently exhibited low power, consistent with its documented limitation arising from additional sub-region testing and the resulting increased multiple-testing burden ^64^. ReMeDy, along with most other methods, maintained reliable FDR control at the nominal 5% level across all sample sizes (Figure 3B), whereas DMRcate exhibited inflated FDR at a sample size of 150. Consistent with these findings, ReMeDy achieved the highest MCC scores across all sample sizes, indicating the most balanced classification of DMRs and non-DMRs (Figure 3C). Similarly, ReMeDy attained the highest AUROC values across all sample sizes, demonstrating superior discrimination between true and non-DMRs (Figure 3D). In contrast, comb-p, DMRcate, and bumphunter showed AUROC values near 0.5 at the smallest sample size (N = 50), with improvement at larger sample sizes but remaining consistently below ReMeDy, while dmrff performed no better than random guessing. ReMeDy’s favorable performance was preserved under equal group sizes, and across other mean effect settings, demonstrating its ability to capture a variety of mean effects and robustness to group imbalance (SFigure 1-5).

**Figure 3:**
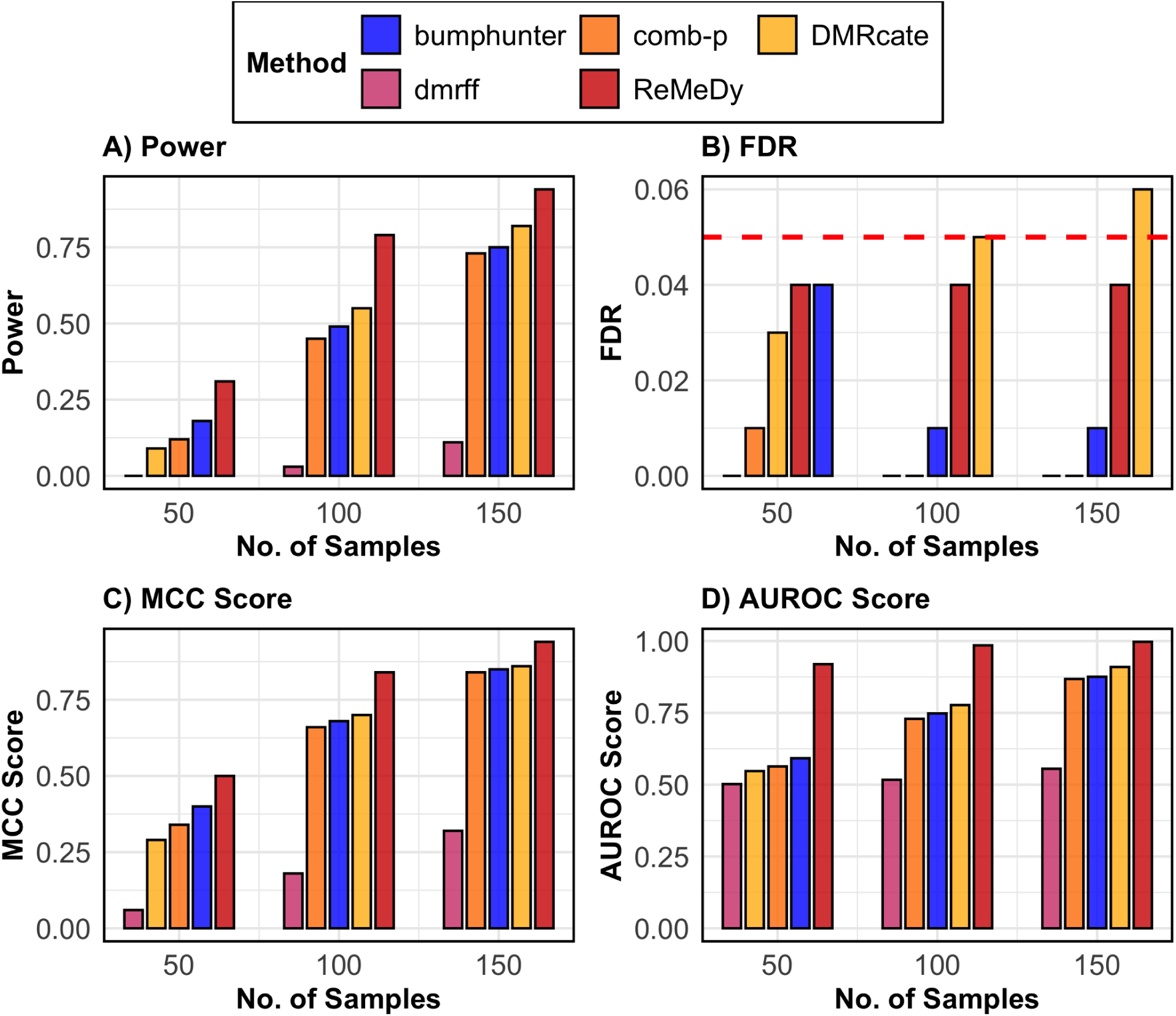
**Performance of ReMeDy and competing methods in the DMR only scenario**. This panel plot compares ReMeDy with other competing methods in DMR only scenario across three sample sizes (50, 100, and 150) with mean effect of 0.7 and unequal group proportions. Panel A shows statistical power, Panel B shows FDR, Panel C shows the MCC score and Panel D shows AUROC scores. Each bar reflects median of 100 simulation runs. The dashed red line in Panel B marks the nominal FDR threshold of 0.05. Higher values in Panels A, C, and D indicate better performance. Methods without a visible bar achieved value of zero.

Building on the DMR-only simulation results, we next evaluated performance under a VMR-only scenario using the same simulation framework of 10,000 regions and 10% true signals, but with no mean effect and a variance in group 2 that was 2.5 times larger than in group 1 (Figure 4). Across all metrics, ReMeDy again demonstrated favorable performance. In terms of power, ReMeDy showed the highest ability to detect true VMRs, with power increasing consistently with sample size (Figure 4A). JLSsc, despite being a site-level method, performed comparably to ReMeDy, while DiffVar and iEVORA showed moderate power that improved with sample size. The comparatively weaker performance of iEVORA reflects its two-step design, in which detection of variability is followed by filtering based on mean differences, resulting in limited signal when only variance effects are present. The VMR detection implementation of DMRcate showed little to no improvement in power across sample sizes, while Bartlett’s test consistently exhibited the lowest power. ReMeDy along with JLSsc, iEVORA, Bartlett’s test and Levene’s test maintained FDR control near the nominal 5% level across all sample sizes (Figure 4B), whereas DMRcate displayed inflated FDR across all sample sizes. In terms of MCC, JLSsc achieved the highest values at some sample sizes (N = 100 and 150), with ReMeDy remaining a close second (Figure 4C). A similar pattern was observed for AUROC, where ReMeDy and JLSsc consistently showed the strongest discrimination between true and non-VMRs (Figure 4D). These results were robust to equal group sizes, with ReMeDy again performing favorably across metrics and across settings with varying variance effects (SFigure 6-10).

**Figure 4:**
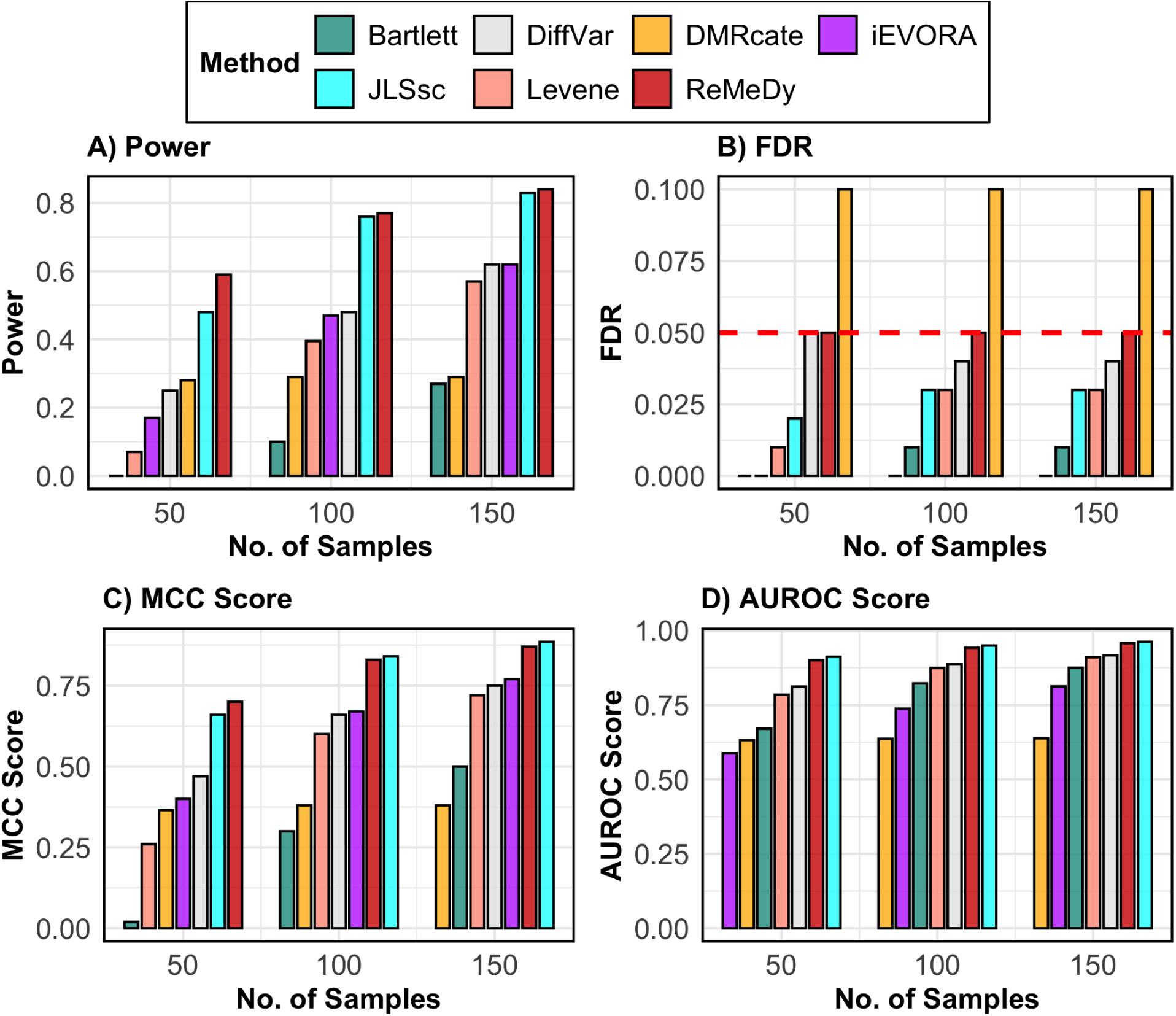
**Performance of ReMeDy and competing methods in the VMR only scenario**. This panel plot compares ReMeDy with other competing methods in VMR only scenario across three sample sizes (50, 100, and 150) with variance effect of 2.5 and unequal group proportions. Panel A shows statistical power, Panel B shows FDR, Panel C shows the MCC score and Panel D shows AUROC scores. Each bar reflects median of 100 simulation runs. The dashed red line in Panel B marks the nominal FDR threshold of 0.05. Higher values in Panels A, C, and D indicate better performance. Methods without a visible bar achieved value of zero.

**Figure 5:**
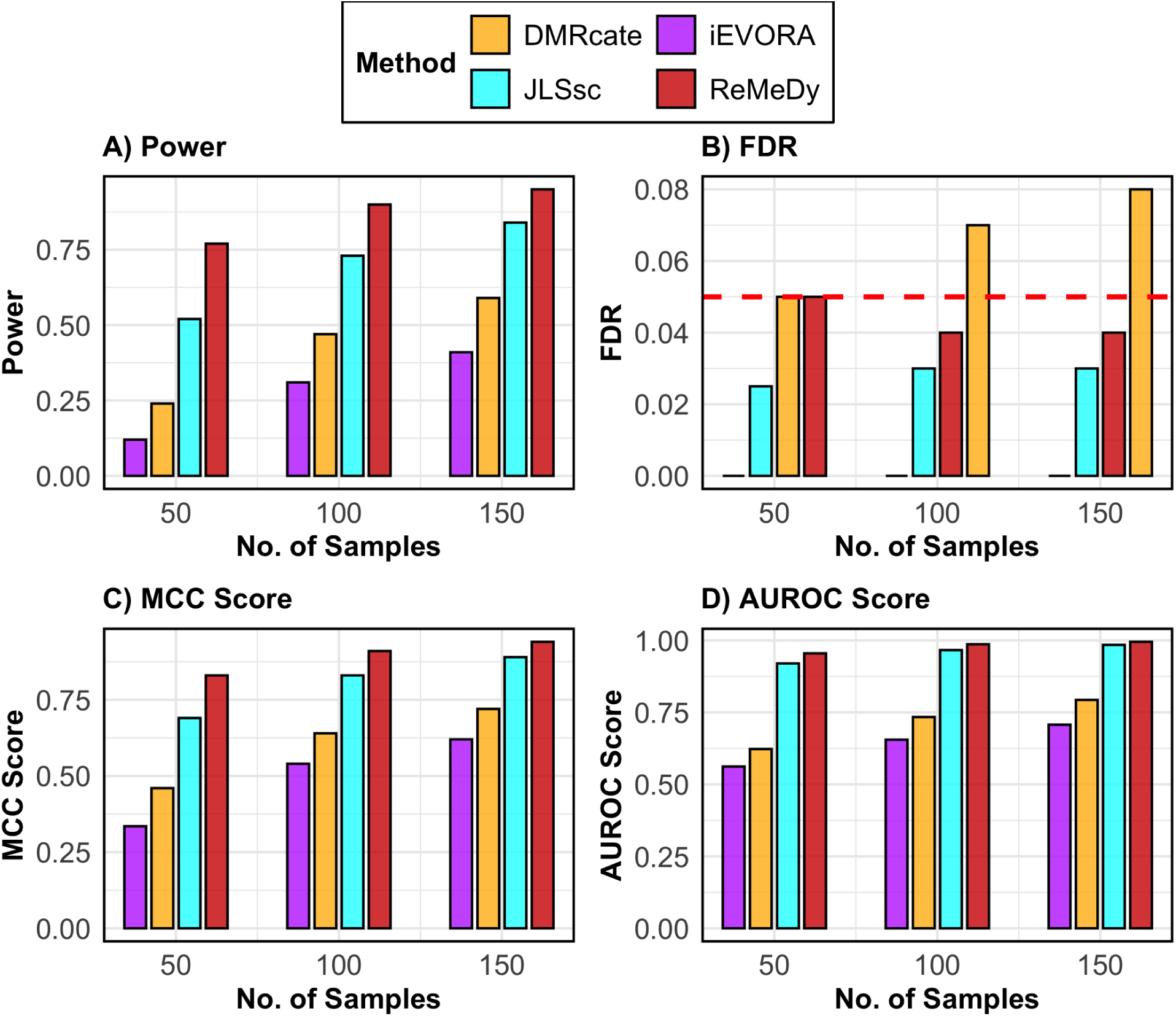
**Performance of ReMeDy and competing methods in the DVMR scenario**. This panel plot compares ReMeDy with other competing methods in DVMR scenario across three sample sizes (50, 100, and 150) with mean effect of 0.7, variance effect of 2.5 and unequal group proportions. Panel A shows statistical power, Panel B shows FDR, Panel C shows the MCC score and Panel D shows AUROC scores. Each bar reflects median of 100 simulation runs. The dashed red line in Panel B marks the nominal FDR threshold of 0.05. Higher values in Panels A, C, and D indicate better performance. Methods without a visible bar achieved value of zero.

**Figure 6:**
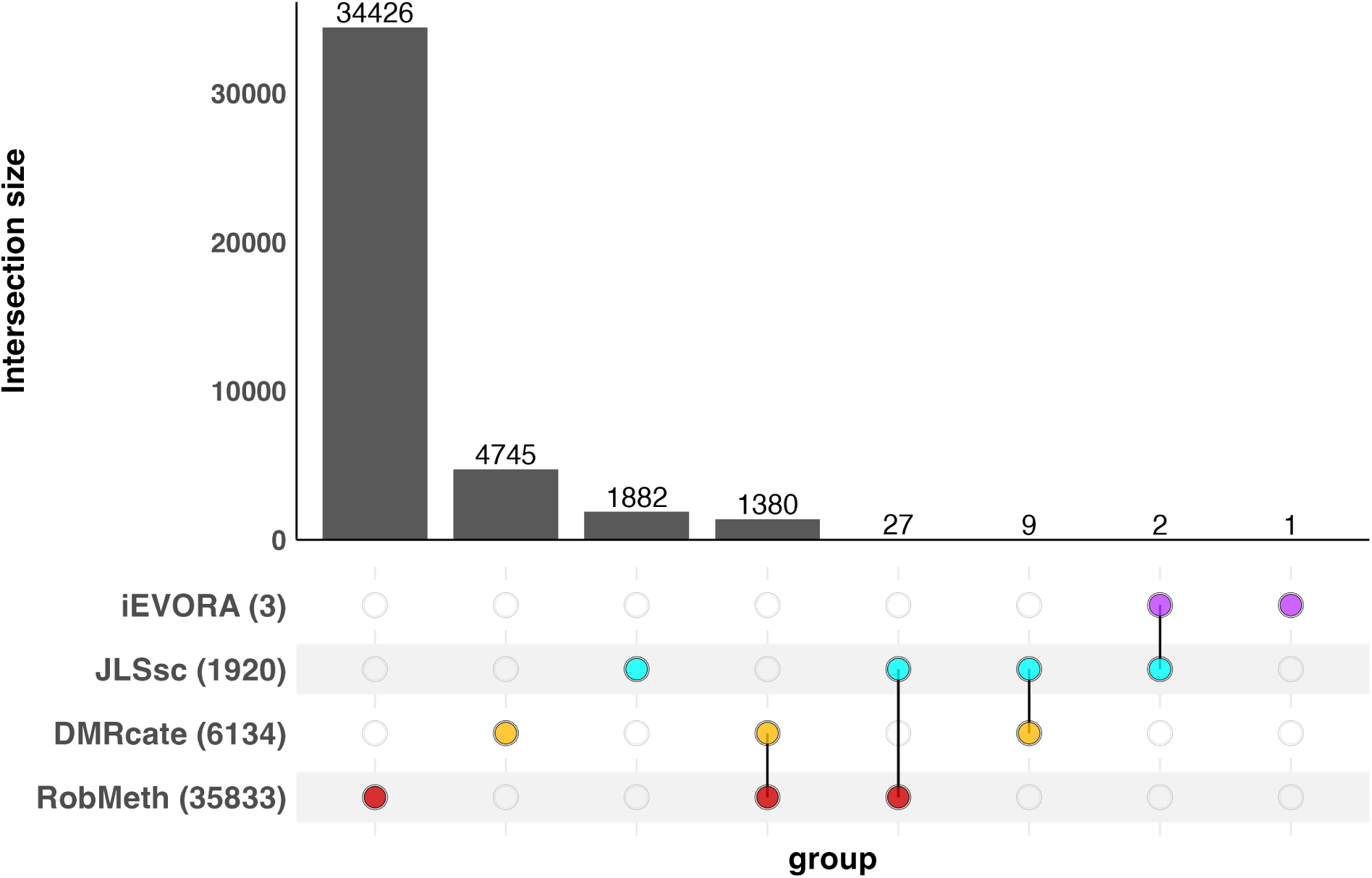
Upset plot of CpG-level overlap across four methylation analysis methods in the pediatric B-ALL dataset. Using four methods, we summarize the overlap of CpGs within the regions identified as differentially methylated, variably methylated or both between B-ALL patients (N = 53) and non-leukemia controls (N = 4) in the pediatric B-ALL dataset generated using the Illumina EPIC array. Numbers in parentheses indicate the total number of CpGs detected by each method. Intersections are evaluated at the CpG level to allow fair comparison between region-based and site-based approaches. CpGs with adjusted p-values smaller than *α* = 0.05 were considered significant by each method.

Finally, in a similar vein, we assessed model performances under the DVMR scenario which incorporated both a mean difference of 0.7 and a variance effect of 2.5, while retaining the same simulation design of 10,000 regions, 10% true signals, and imbalanced group proportions (Figure 5). Across all evaluation metrics, ReMeDy continued to demonstrate superior performance. ReMeDy achieved the highest statistical power across all sample sizes, with steady improvement as sample size increased (Figure 5A). JLSsc and DMRcate also showed increasing power, while iEVORA exhibited limited improvement, reflecting again the inefficiency arising from its sequential mean-filtering step when both effects coexist. Similar to our previous findings,

ReMeDy along with JLSsc and iEVORA consistently maintained FDR near the nominal 5% level across all sample sizes (Figure 5B), whereas DMRcate showed inflated FDR at moderate and large sample sizes. MCC and AUROC analyses further highlighted ReMeDy’s favorable performance, with ReMeDy achieving the highest scores across sample sizes and JLSsc consistently ranking second (Figure 5C–D) while DMRcate and iEVORA demonstrated only moderate discrimination capabilities, indicating limited adaptability to combined mean–variance effects. Consistent performance was observed under equal group sizes, with ReMeDy maintaining favorable results across evaluation metrics and under varying mean and variance effects (SFigure 11-27).

### 3.4 Re-analysis of population-level DNAm array dataset

We used ReMeDy to analyze a publicly available DNAm array dataset from the Gene Expression Omnibus database (GEO). The dataset (GSE306095) was generated using the Illumina EPIC platform, and quality control and batch effect correction were performed using the ChAMP ^63^ pipeline and ComBat ^61^ (See supplementary section 1.3 for more details). This dataset investigates DNAm changes in B-cell acute lymphoblastic leukemia (B-ALL) and comprises of 57 samples, including 53 B-ALL patients and 4 non-leukemia controls, with methylation measured at 719,234 CpG sites ^69^. Childhood ALL is a blood and bone marrow malignancy characterized by uncontrolled proliferation of immature lymphoid cells and represents the most common pediatric cancer, with symptoms including fatigue, bruising, fever, bone pain, and recurrent infections ^70^. Standard treatment involves multi-phase chemotherapy, often combined with targeted or immunotherapy, resulting in high long-term survival rates ^71;72^. We selected this dataset because it represents a very recent EWAS that focused exclusively on identifying mean methylation differences associated with B-ALL, thereby overlooking other important components of methylation dysregulation, such as differential variability and joint mean–variance effects ^69^. The original study reported a predominantly hypermethylated profile in B-ALL and identified 1,056 DMRs enriched in pathways related to cell proliferation and neuronal signaling. Strong associations were also observed at CpGs within *MAD1L1* and *RPTOR*. However, this analysis primarily focused on mean differences, providing an incomplete view of the B-ALL associated methylation landscape ^69^. By applying ReMeDy, we aim to recover additional variance-driven and joint mean–variance methylation signals that may further refine the epigenetic characterization of B-ALL and reveal complementary disease-associated patterns not captured by the current standard of EWAS.

In addition to analyzing the B-ALL dataset with ReMeDy, we performed comparative analyses using DMRcate, JLSsc, and iEVORA, as these methods can also identify all three components of methylation dysregulation. ReMeDy identified the largest number of CpGs in the B-ALL dataset (35,833 CpGs grouped into 7,246 regions), compared with DMRcate (6,134 CpGs), JLSsc (1,920 CpGs), and iEVORA (3 CpGs) (Figure 6). Among the regions detected by ReMeDy, 383 were classified as DMRs (1,680 CpGs), 3 as VMRs (12 CpGs), and 6,860 as DVMRs (34,141 CpGs). To ensure fair comparison across methods that output either individual CpGs or regions containing multiple CpGs, overlaps were assessed at the CpG level. ReMeDy and DMRcate shared 1,380 CpGs, corresponding to an overlap of approximately 22.5%, indicating concordance between the two region-based approaches. Notably, ReMeDy uniquely identified 34,426 CpGs across 7,000 regions, the majority of which were DVMRs (6,622 regions comprising 32,774 CpGs), along with additional unique DMRs (375 regions, 1,640 CpGs) and VMRs (3 regions, 12 CpGs). Overall, these results demonstrate that ReMeDy achieves both agreement with established methods and, consistent with the simulation studies, exhibits substantially broader detection power by capturing additional methylation signals that are not identified by existing approaches.

Following data analysis, we performed functional bioinformatics analyses to assess the biological relevance of CpGs located in regions uniquely identified by ReMeDy (See supplementary section 1.4 for more details). CpG island annotation showed that a substantial proportion of these CpGs mapped to CpG islands (55.2%), with additional enrichment in open sea (20.2%) and shore regions, indicating that ReMeDy captures both promoter-proximal and distal regulatory methylation changes (Figure 7B). Genic annotation further revealed that most CpGs were located within gene-associated regions, including promoters (34.1%), regions 1–5 kb upstream of transcription start sites (24.7%), introns (26.2%), and exons (10.1%), highlighting their potential regulatory impact on gene expression (Figure 7C). Among regions uniquely identified by ReMeDy, 18.3% exhibited low effect sizes, while the majority (81.7%) showed moderate to high effect sizes, indicating sensitivity to both subtle and strong methylation changes. The top 10 significant regions and their corresponding effect sizes are shown in Figure 7D. Pathway-level analyses identified enrichment of transmembrane receptor protein tyrosine phosphatase signaling among the regions uniquely identified by ReMeDy (Figure 7A), driven in part by *PTPRD*, a receptor-type protein tyrosine phosphatase frequently disrupted or deleted in pediatric and adult ALL, including at relapse ^73;74^. Given the central role of receptor-type protein tyrosine phosphatases in intracellular signal transduction, whose loss promotes aberrant signaling and leukemic progression ^75^, these results support the biological plausibility of regions uniquely identified by ReMeDy. Consistent with this, Gene Ontology (GO) ^76^ enrichment analysis further highlighted Wnt signaling as a key pathway associated with regions uniquely identified by ReMeDy (Figure 7E), with multiple genes (*WNT3A*, *WNT7A*, *ROR2*, *FZD10*, *CSNK1E*) involved in canonical and non-canonical Wnt cascades. Dysregulation of Wnt signaling is known to influence hematopoietic stem cell self-renewal, leukemic cell survival, aberrant *β*-catenin activation, and treatment resistance in ALL ^77;78^. Together, these findings indicate that ReMeDy captures methylation dysregulation linked to biologically meaningful pathways central to disease pathogenesis.

**Figure 7:**
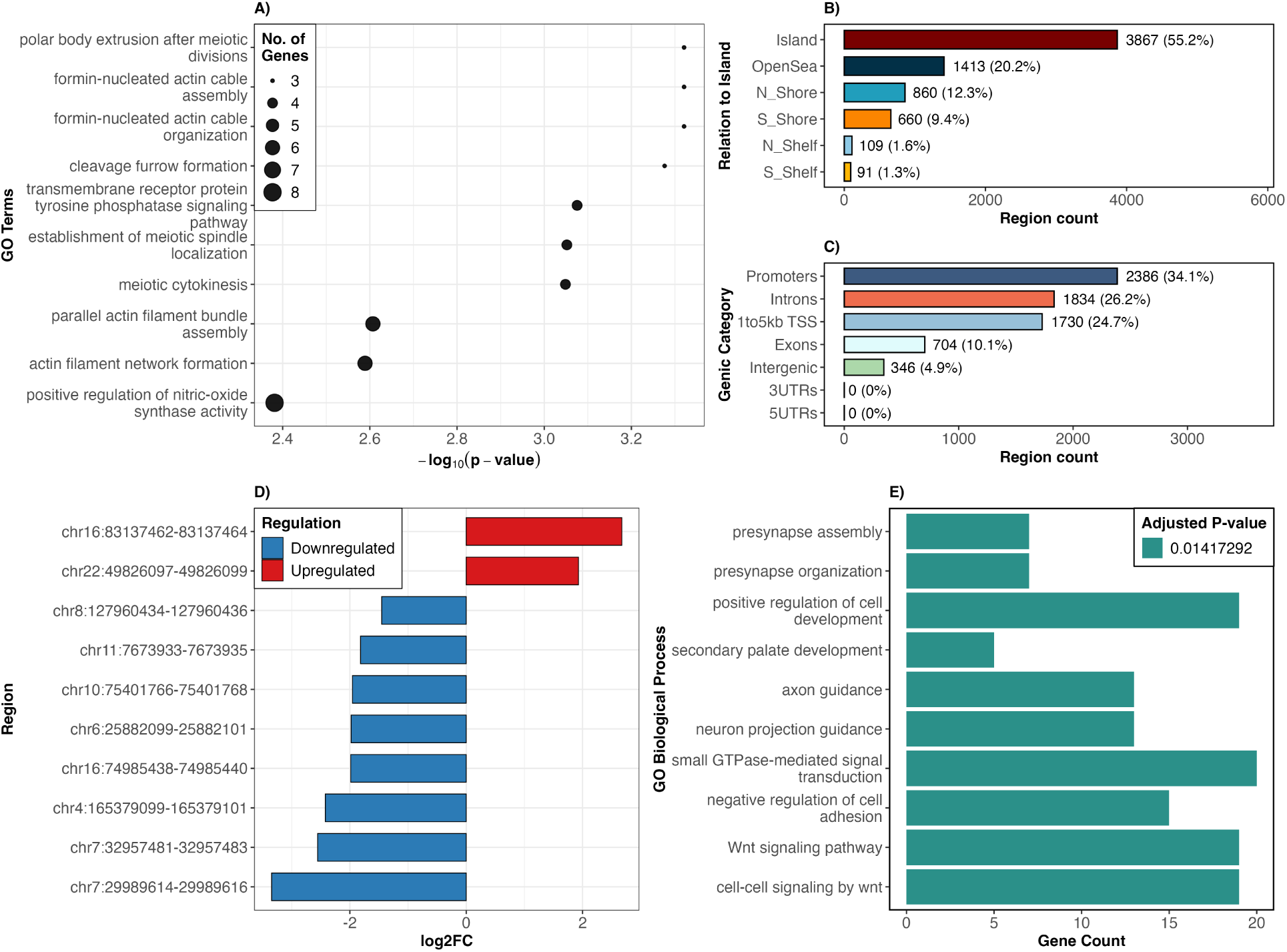
This figure summarizes the genomic context, functional annotation and effect sizes of regions uniquely identified by ReMeDy. Panel A shows GO biological process enrichment derived from region-based analysis using goregion() ^88^, where genomic regions were directly used to identify enriched functional terms; point size indicates the number of genes associated with each term and the x-axis shows −*log*_10_(*p* − *values*). Panel B shows the distribution of these regions relative to CpG islands, including islands, shores, shelves and open sea regions. Panel C shows the genic annotation of CpGs within these regions, categorized by promoters, regions 1-5 kb upstream of transcription start sites, exons, introns, untranslated regions and intergenic regions. Panel D shows the log2FC of the top 10 regions ranked by effect size, with positive and negative values indicating upregulated and downregulated regions, respectively. Panel E shows GO biological process enrichment based on genes associated with ReMeDy-specific regions, obtained using gene-level enrichment analysis (enrichGO ^89^), with bar lengths representing the number of genes contributing to each pathway.

We further mapped CpGs in regions uniquely identified by ReMeDy to their nearest genes, revealing several biologically relevant genes associated with B-ALL. Among the 7,000 regions uniquely identified by ReMeDy, one of the strongest signals (chr22:49826097–49826265) mapped to *BRD1*, an epigenetic regulator reported as a rare fusion partner of *PAX5* in childhood B-cell precursor ALL, supporting its functional relevance in leukemogenesis ^79;80^. Two additional regions on chromosome 21 (chr21:34886125–34886300 and chr21:34886769–34887501) mapped to *RUNX1*, a key hematopoietic transcription factor frequently involved in childhood B-ALL through the *ETV6::RUNX1* fusion, which drives leukemic transformation in a major clinical subtype ^81^. Recent studies have also reported rare pathogenic *RUNX1* point mutations in relapsed B-ALL, including a novel p.Leu148Gln variant associated with poor clinical outcome ^82^. Another region (chr21:38499362–38499580) mapped to *ERG*, an ETS-family transcription factor implicated in a distinct subtype of B-cell precursor ALL. Recurrent *ERG* deletions occur predominantly in *DUX4* -rearranged B-ALL, where *DUX4* deregulation promotes expression of a dominant-negative *ERG* isoform or predisposes the locus to deletion, contributing to leukemogenesis and defining a biologically distinct subgroup ^83;84^. The presence of these well-established B-ALL associated genes among regions uniquely detected by ReMeDy supports its ability to identify biologically meaningful epigenetic signals. At the same time, a subset of regions uniquely identified ReMeDy mapped to genes with no currently known links to B-ALL, suggesting the potential discovery of novel regulatory elements or candidate biomarkers that warrant further functional and clinical investigation.

## 4 Discussion

In this study, we introduced ReMeDy, an integrated statistical framework for region-based identification of methylation dysregulation, encompassing differential methylation, differential variability, and their joint effects. Methodologically, ReMeDy is built on a hierarchical generalized linear modeling framework that naturally incorporates correlation among CpG sites and enables joint mean–variance modeling with respect to the phenotype of interest and covariates under one statistical framework, thereby addressing key shortcomings of current EWAS pipelines (Table 1). With a computationally efficient hierarchical likelihood–based estimation framework and region-wise model fitting, ReMeDy is scalable to DNAm datasets comprising tens of thousands of genomic regions measured across large sample cohorts. While the primary use case presented here focuses on univariate analysis with a binary phenotype, extension to continuous or discrete phenotypes, multivariable modeling with multiple covariates or confounders, and incorporation of additional random effects is straightforward. Beyond array-based methylation data, the ReMeDy framework is readily extensible to sequencing-based methylation data and other high-dimensional modalities with correlated structure, including histone modification profiles, ATAC-seq, copy number variation, and microRNA expression data. Collectively, these features position ReMeDy as a flexible and powerful statistical framework that expands the scope of methylation dysregulation detectable in EWAS and other related genomic analyses.

**Table 1:**
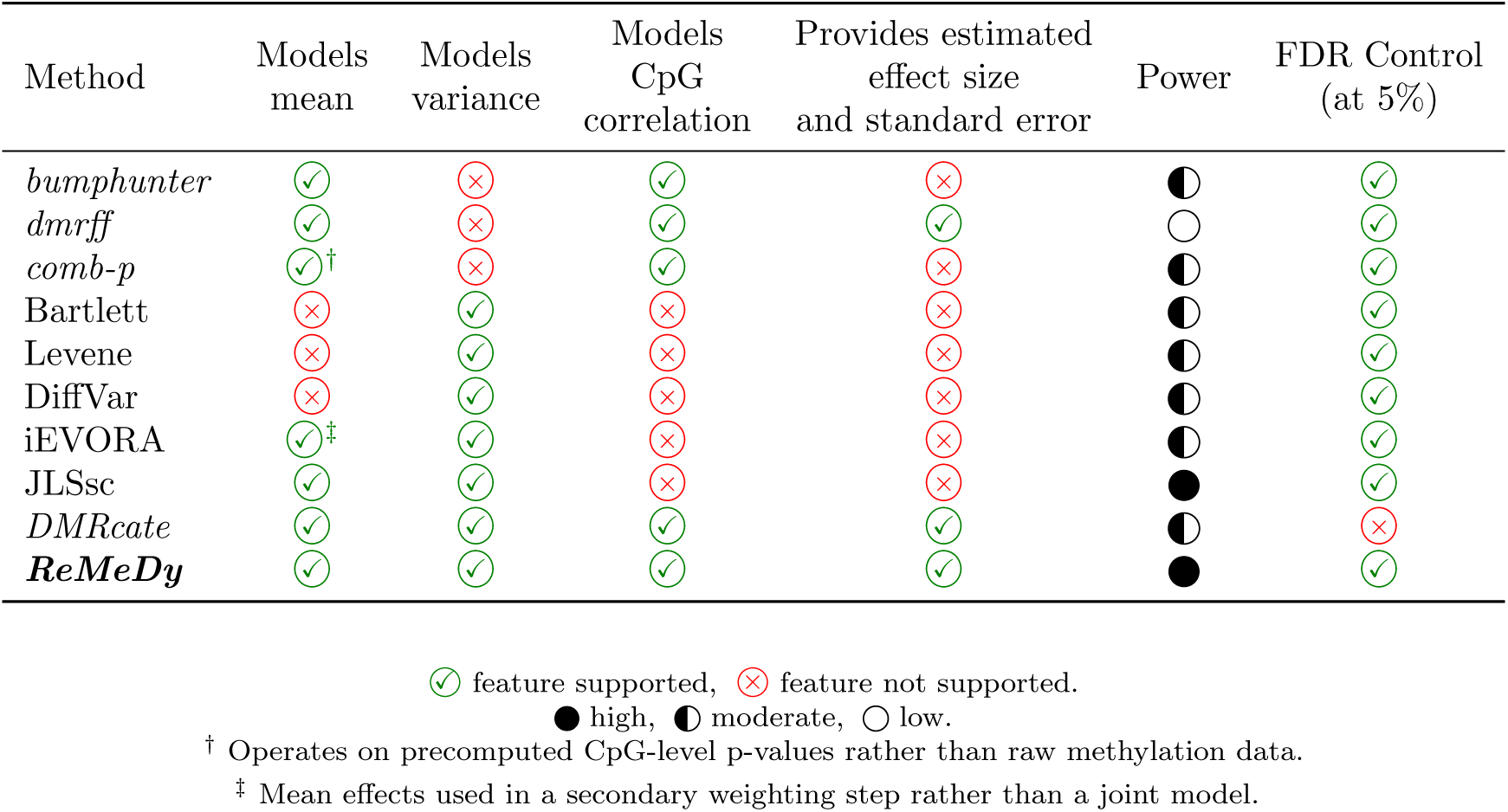
This table summarizes key methodological characteristics and performance of ReMeDy and other methods for identifying methylation dysregulation. The first column lists the methods. The second and third columns indicate whether each method explicitly models mean and variance effects, respectively. The fourth column specifies whether CpG-level correlation within regions is accounted for. The fifth column indicates whether the method provides estimated effect sizes and associated standard errors. The sixth column summarizes empirical power, and the final column reports FDR control at the nominal 5% level. The methods in italics perform region-based analysis.

Given the substantial body of existing methodological work in this area, it is important to clearly articulate the practical strengths and implementation advantages of ReMeDy. First, ReMeDy operates directly on biologically defined co-methylated regions, allowing it to naturally account for the inherent correlation structure present in DNAm array data, rather than reconstructing regions post hoc from site-level association statistics using smoothing, clustering, or p-value combination approaches. Second, unlike many existing region-based methods, ReMeDy does not rely on heuristic tuning parameters such as smoothing spans, kernel bandwidths, autocorrelation windows, or CpG-level significance thresholds that can substantially influence results and introduce subjectivity ^65;38;48;64^. For example, DMRcate requires users to specify kernel bandwidth (*λ*) and scaling (*C*) parameters, where different choices can lead to markedly different region sizes and numbers of discoveries. In contrast, ReMeDy performs inference within a single, coherent statistical model, yielding results that are robust to arbitrary user-defined settings. Third, ReMeDy achieves computational efficiency by replacing full marginal likelihood evaluation with hierarchical likelihood optimization, thereby avoiding high-dimensional numerical integration and enabling stable and efficient parameter estimation without the need for repeated resampling or intensive permutation procedures (SFigure 28). Fourth, ReMeDy can incorporate multiple random effects, allowing it to model subject-level variation alongside additional sources of correlation, such as tissue-specific effects ^85^, thereby making it well suited for emerging multi-tissue methylation studies ^86;87^. Finally, ReMeDy has been implemented in R in a parallelizable manner and is freely available through our GitHub repository, facilitating scalable computation, accessibility, reproducibility, and future methodological development by the broader research community.

The extensive null and power simulations conducted in this study provide strong empirical support for the statistical validity of ReMeDy. Under null scenarios, ReMeDy consistently maintained type-I error rates close to the nominal level across a range of sample sizes, demonstrating well-calibrated inference despite the added complexity of joint mean–variance modeling and region-level correlation structures. In contrast, several existing methods exhibited either excessive conservatism or mild inflation, particularly in small-sample settings, highlighting the challenges of error control in spatially correlated methylation data ^40;39^. Across power simulations, ReMeDy showed robust and often superior performance in detecting true differential, variable, and joint mean–variance methylation signals, with gains becoming more pronounced as sample size increased. Notably, ReMeDy retained strong performance in scenarios driven purely by variance effects or by combined mean–variance changes, settings in which many competing approaches showed limited sensitivity due to their reliance on mean-only or sequential testing strategies. These results demonstrate that ReMeDy achieves a favorable balance between statistical power and false discovery control across a broad range of realistic EWAS conditions.

Application of ReMeDy to a analyze a recent pediatric ALL data demonstrated the practical value of moving beyond mean-centric analyses. While the original study focused exclusively on DM, ReMeDy identified a substantially broader set of ALL-associated genomic regions, the majority of which exhibited joint mean–variance dysregulation. Functional annotation showed that genomic regions uniquely identified by ReMeDy but missed by existing models were enriched in regulatory genomic contexts and displayed moderate-to-large effect sizes, supporting their biological relevance. Pathway enrichment analyses further identified signaling processes, including receptor-type protein tyrosine phosphatase and Wnt signaling, which are well established in B-ALL biology but were not fully captured by mean-based or variance-based models alone. Consistent with these findings, gene-level mapping revealed well-characterized B-ALL-associated genes such as BRD1, RUNX1, and ERG, reinforcing the biological plausibility of the detected signals. At the same time, a subset of regions mapped to loci without known links to B-ALL, highlighting ReMeDy’s potential to identify novel regulatory elements and candidate biomarkers. Although further experimental validation is required to establish functional relevance, the ability of ReMeDy to recover both known and previously uncharacterized signals underscores its utility for hypothesis generation and illustrates how integrated region-level joint mean–variance modeling enables a more complete and nuanced characterization of methylation dysregulation in complex diseases.

## Availability of Data and Materials

The 450K datasets are accessible from the GEO database under accession number GSE44667 and GSE80970. The EPIC dataset is accessible from the GEO database under accession number GSE306095. Code for ReMeDy, simulations, and comparisons of methods identifying methylation dysregulation on the various datasets explored are available on GitHub at https://github.com/SChatLab/ReMeDy. The source code is licensed under an MIT License.

## Acknowledgments

This research was supported in part by Lilly Endowment, Inc., through its support for the Indiana University Pervasive Technology Institute.

