## Supplementary Materials for "ReMeDy: A Flexible Statistical Framework For Region-based Detection of DNA Methylation Dysregulation"

**2 Supplementary Figures** **7**

**3 Supplementary Tables** **35**

### 1 Supplementary Text

#### 1.1 Simulation Framework

##### 1.1.1 Power Simulations

To evaluate methods for detecting methylation dysregulation, we generated synthetic DNA methylation (DNAm) datasets that resemble the structure of Illumina 450K array data. A population-level methylation data from Gene Expression Omnibus (GEO; `https://www.ncbi.nlm.nih.gov/geo/`) database under the accession number GSE44667 was used solely to anchor the simulations in a biologically realistic genomic context; further details on this dataset are provided in Section 1.3. The methylation beta matrix was loaded along with the platform manifest, which was filtered to retain only probes present in both sources. Genomic regions were then defined by grouping CpG sites that fell within 200 bp of one another, and each region was required to contain exactly three CpGs. Regions not meeting this criterion were discarded, producing a catalog of candidate regions with genomic coordinates and CpGs assigned to each region ordered by chromosome. The methylation values themselves were not derived from the real dataset. Instead, CpG names extracted from each region served as the row identifiers for the simulated data so that downstream region-finding methods, which require valid probe IDs, could operate. The simulation framework then generated synthetic M-values across a user-specified number of regions, assigning each region a fixed number of CpGs that inherited genomic identifiers from the template. To mimic realistic co-methylation behavior, each region was assigned a correlation structure. A subset of regions was designated as high-correlation regions, while the remainder formed a low-correlation set. For each region, an intra-region correlation value was drawn from a predefined range and a correlation matrix was constructed under a compound symmetric structure. This matrix defined the dependence among CpGs during multivariate sampling. Samples were split into two exposure groups to introduce group structure. Depending on the simulation setting, regions were labeled as differentially methylated (DMRs), variably methylated (VMRs) or both (DVMRs). DMRs and DVMRs received a mean shift in group 2, while VMRs and DVMRs received group-specific variances drawn from scaled inverse-chi-square distributions. Correlated M-values were then generated for each region, and the group-specific variances were applied after multivariate sampling to preserve the intended correlation pattern. All simulated CpGs were annotated with their region, effect type (DMR, VMR, DVMR), mean and variance effects, and the intended correlation used for that region. Empirical correlations were also computed for quality checks. The final dataset consisted of synthetic methylation measurements organized by genomic regions, with biologically plausible correlation patterns and clear ground truth labels to support power evaluations.

##### 1.1.2 Null Simulations

To evaluate type I error, we constructed DNAm datasets with no mean and variance effects. We used two Illumina 450K datasets to represent different sample size settings: a smaller dataset with 40 samples and a larger dataset with 142 samples (details on these datasets are provided in Section 1.3). For each dataset, beta values and sample metadata were loaded along with the 450K manifest, which was filtered to retain only probes present in the methylation data. To remove any association between methylation levels and exposure groups we randomly permuted the sample columns of the beta matrix. We then defined genomic regions by grouping CpGs that were within 200 bp of one another and retaining only regions containing exactly three CpGs. From these candidate regions, 10,000 regions were sampled at random. The permuted beta matrix was subset to the CpGs in these regions, samples were relabeled to create two equal-sized exposure groups and beta values were converted to M-values using ENmix::B2M[33]. The resulting datasets preserve realistic probe spacing and within-region correlation while containing no mean or variance differences between groups.

#### 1.2 Details of Statistical Models

In this section, we describe the statistical methods used in our study to identify methylation dysregulation which were evaluated alongside ReMeDy. These included bumphunter[13], comb-p[22], DMRcate[23], dmrff[29], JLSsc[28], iEVORA[30], DiffVar[25], and the Bartlett's[2] and Levene's[17] tests. We also describe SACOMA[20], which was used to identify co-methylated regions that serve as input to our model ReMeDy.

**bumphunter:** bumphunter first applies a linear regression model to each CpG site, considering confounding factors like age, sex, and batch effects. Then, it smooths out these changes across neighboring sites to find 'bumps' or clusters that might represent meaningful differences. These candidate regions are determined by setting a threshold for how big the bumps need to be. The method then uses permutation tests or a faster alternative called bootstrapping to assess how likely these regions are to occur by chance, adjusting for statistical errors to ensure the results are reliable. bumphunter is implemented in various R packages, like bumphunter, ChAMP[21] and minfi[1].

**DMRcate:** DMRcate starts by fitting a linear model to each CpG site, using a method that leverages Bayesian statistics to assess the effect of group status on methylation values. The resulting t-statistics are squared and smoothed using a Gaussian kernel with a specific bandwidth. DMRcate then computes p-values for each CpG using the Satterthwaite[26] method, then performs adjusting for multiple comparisons using the Benjamini-Hochberg method[3] to control for false discoveries. Sites with adjusted p-values below a set threshold (like 0.05) are considered significant, and those close together are grouped into regions. The smallest p-value within each region is used to indicate its significance level. In addition to identifying DMRs, DMRcate also provides functionality for detecting VMRs and DVMRs. This functionality is documented in its package documentation. DMRcate is implemented as a bioconductor package.

**comb-p:** comb-p takes a file containing p-values and chromosome locations of CpG sites as input, instead of calculating p-values for individual CpGs. It first calculates the autocorrelation of p-values at varying distances to understand the correlation between neighboring probes. The tool then applies the Stouffer–Liptak–Kechris (SLK)[11;18] correction to adjust the p-values by considering the autocorrelation and nearby p-values. After adjusting the p-values, comb-p employs a peak-finding algorithm to identify regions enriched with small p-values, indicating potential DMRs. The regional significance is then recalculated using the original p-values with the Stouffer–Liptak correction, along with an additional Sidak correction[35] to account for multiple testing. comb-p is available as a command-line utility, a python library and through R package ENmix.

**dmrff:** dmrff identifies DMRs by combining CpG-level association results from an epigenome-wide association study (EWAS) while explicitly accounting for correlation between nearby CpG sites. It starts from EWAS summary statistics and defines candidate regions as clusters of nearby CpGs with nominal significance and consistent direction of effect. For each candidate region and its sub-regions, dmrff computes a region-level statistic using an extension of inverse-variance weighted meta-analysis that incorporates the CpG correlation structure. The most significant non-overlapping sub-regions are then selected and region-level p-values are adjusted for multiple testing using Bonferroni correction.

**Bartlett's test:** Bartlett's test used to check if multiple groups have equal variances or if at least two of them differ in variance. It is an extension of the F-test and is highly sensitive to outliers and deviations from normality. Moreover, it doesn't allow for adjusting covariates in its analysis.

**Levene's Test:** Levene's test is used to check whether variability is comparable across two or more groups. Compared to Bartlett's test, it is less affected by violations of normality, which makes it more reliable in many real-world settings. The test works by calculating how far each observation deviates from its group mean or, in a more robust version, the median. These deviations are then analyzed using an ANOVA framework. If the variances are equal across groups, the test statistic follows an F distribution. A statistically significant result suggests that at least one group differs in variance, and the assumption of equal variances is rejected.

**DiffVar:** DiffVar is a method for detecting differences in variability between groups in high-dimensional DNAm data. Building on the method of Levene's test, it evaluates variability by examining how far each observation lies from its group average, which makes it possible to identify features with group-specific differences. When testing across thousands of CpG sites, DiffVar reduces the risk of false positives by using moderated t-statistics. The method is implemented within a linear modeling framework, allowing for covariate adjustment and accommodating complex experimental designs, while remaining robust to outliers.

DiffVar is available as part of the missMethyl package in R.

**iEVORA:** iEVORA (improved Epigenetic Variable Outliers for Risk Prediction Analysis) is a regularized extension of the EVORA method. It begins by using Bartlett's test to identify CpG sites where methylation variability differs between groups. Because Bartlett's test can be highly sensitive to outliers, iEVORA refines this initial set by re-ranking the identified sites, placing greater emphasis on those that also show meaningful differences in average methylation levels. This approach yields a more refined set of markers that capture regions where both variable and differential methylation changes are most pronounced.

**JLSsc:** The Joint Location and Scale Score (JLSsc) test brings together two complementary analyses, one focused on differences in group means (location) and the other on differences in variability (scale). By accounting for the correlation between these two components, JLSsc offers a more flexible and robust framework. It can be applied to both continuous and categorical variables and allows for adjustment of covariates. The method is implemented in the jlst R package through the jlssc function, which gives users the option to assess variance differences using one of four approaches: the Breusch–Pagan test [7], Brown–Forsythe test [8] or method of moments versions of either test.

**SACOMA:** SACOMA is a data-driven, unsupervised method for identifying co-methylated regions by clustering neighboring CpG sites that show correlated methylation patterns. It models region detection as a spatially constrained hierarchical clustering problem, jointly accounting for methylation similarity and genomic proximity. SACOMA uses a data-adaptive mixing parameter which helps in avoiding rigid assumptions. CpG sites are first grouped into local genomic bins, and within each bin, correlated CpGs are clustered to define co-methylated regions. This approach reduces dimensionality while preserving biologically meaningful methylation structure and provides a robust foundation for downstream region-based analyses.

#### 1.3 Data Description

In this study, we obtained three publicly available methylation dataset from the GEO database under the accession numbers GSE44667 (EOPET dataset), GSE80970 (Alzheimer dataset) (both based on the 450K platform) and GSE306095 (B-ALL dataset, based on EPIC platform). Both the 450K datasets were used in the type-I error evaluation and the EPIC dataset was used for population-level analysis in our study.

The EOPET dataset focuses on DNAm patterns in placental tissue from cases of early-onset preeclampsia (EOPET) [6]. It includes 40 samples in total, with 20 EOPET and 20 control samples. Methylation profiling was performed using the Illumina Infinium HumanMethylation450 BeadChip, which measures over 480,000 CpG sites genome-wide [6]. During preprocessing, probes located on sex chromosomes and those containing known single-nucleotide polymorphisms at the C or G positions were removed. Probes with detection p-values greater than 0.01 and probes with missing beta values were also excluded, resulting in a final set of 430,685 probes. Signal intensities were imported into R using the methylumi [10] package and converted to M-values. The data were then normalized using subset-quantile within-array normalization [19], after which the values were transformed back to beta values [6].

The Alzheimer dataset investigates changes in DNAm in prefrontal cortex and superior temporal gyrus samples from 147 individuals [27]. In our study we used the samples from the prefrontal cortex, which comprises 142 samples, including 74 from patients with Alzheimer's disease and 68 controls [27]. The dataset was profiled using Illumina's Infinium HumanMethylation450 BeadChip, which assesses over 480,000 CpG sites across the genome. For initial data quality control, the GenomeStudio (version 2011.1) was used to assess key metrics, including staining, extension, hybridization, target removal, bisulfite conversion efficiency, specificity, and nonpolymorphic and negative controls. Probes known to map to multiple genomic locations or containing single-nucleotide polymorphisms (SNP) at the single-base extension site were excluded from further analysis, along with the 65 SNP probes used for sample identification on the array, resulting in the removal of 72,067 probes in total. DNAm levels for each probe were quantified using beta values, defined as the ratio of the methylated signal to the total signal intensity (M / [M + U]) [27].

The B-ALL dataset was generated using the Illumina EPIC platform, which investigates DNAm changes in B-cell acute lymphoblastic leukemia (B-ALL)[12]. This dataset comprises of 57 samples, including 53 B-ALL patients and 4 non-leukemia controls[12]. The initial quality control of the raw IDAT files was carried out using the BeadArray Controls Reporter (Illumina, San Diego, CA, USA). Data preprocessing and normalization were performed in R (v4.3.0) with the ChAMP[21] pipeline (v2.23.0) for EPIC arrays, obtained from Bioconductor v3.18. Probes with detection p-values $\geq 0.01$ in more than 5% of samples were removed due to low signal quality. We further excluded non-CpG probes, probes containing single-nucleotide polymorphisms at the CpG site or single-base extension, multi-hit probes mapping to multiple genomic locations, and probes on the X and Y chromosomes to minimize sex-related confounding. Samples in which more than 10% of probes failed the detection p-value threshold were also excluded. To correct for probe design bias between type I and type II probes, intra-array normalization was performed using the Beta Mixture Quantile method[31]. Technical batch effects were addressed using the ComBat[14] algorithm from the sva[16] R package, with sample plate included as a known batch variable. DNAm levels were quantified as beta values, defined as the ratio of methylated signal intensity to the total signal intensity at each CpG site, ranging from 0 to 1. All probes were mapped and annotated using the GRCh37/hg19 human genome reference[12].

#### 1.4 CpG, Genic and Functional Annotation

For CpG annotation, significant regions with methylation dysregulation identified by our method were first extracted, and the CpGs within these regions were linked to the appropriate array manifest file (HM450 or EPICv). Regional annotations were summarized by determining the most frequent CpG neighborhood category within each region, including Island, Shore, Shelf, and OpenSea. Genic annotation was performed in R using the annotatr[9], GenomicRanges[15], and TxDb.Hsapiens.UCSC[4;5] packages. For each method, CpG sites of each regions were converted to genomic coordinates and annotated against gene-based features derived from the corresponding TxDb object and annotatr gene models. CpGs located 1–5 kb upstream of transcription start sites were identified separately using promoter definitions from GenomicRanges. Each CpG was assigned to a functional category-Promoters, 1–5 kb TSS, Exons, Introns, 5'UTRs, 3'UTRs, or Intergenic, according to a predefined hierarchical priority rules. Region-level annotations were then determined by assigning each region the predominant genic category represented among its CpGs.

To assess the functional relevance of the identified regions, pathway enrichment analyses were performed using Gene Ontology (GO)[32] biological process terms. Region-based enrichment was conducted using the goregion() function from the missMethyl[24] R package, which directly tests genomic regions for overrepresented functional annotations. In parallel, gene-based enrichment analysis was performed by mapping significant regions to their associated genes and evaluating GO biological process enrichment using the enrichGO() function of the clusterProfiler[34] R package.

#### 1.5 Evaluation Metrics

For evaluating performance of ReMeDy and all the other methods in detecting methylation dysregulation we performed comprehensive comparison using key metrics like Type-I error, false discovery rate (FDR), statistical power, area under the ROC curve (AUROC score) and Matthews correlation coefficient (MCC score).

**Type 1 Error:** The probability of incorrectly rejecting a true null hypothesis. It represents the risk of falsely identifying significant difference when there is none.

$$\text{Type 1 Error} = \frac{\text{FP}}{\text{FP} + \text{TN}}$$

**FDR:** FDR represents the expected proportion of false positives among all significant findings.

$$\text{FDR} = \frac{\text{FP}}{\text{FP} + \text{TP}}$$

**Statistical Power:** It represents the likelihood that the hypothesis test will correctly reject the null hypothesis.

**Mathew's Correlation Coefficient (MCC score):** It evaluates the quality of binary and multi-class classifications by providing a single value that summarizes the information in confusion matrix. It ranges from -1 to 1: a score of 1 indicates perfect predictions, 0 indicates completely random predictions, and -1 indicates predictions that are completely opposite of the actual outcomes. MCC evaluates models based on true positives, true negatives, false positives, and false negatives, helping to minimize significant errors in findings.

$$\text{MCC} = \frac{\text{(TP * TN) - (FP * FN)}}{\sqrt{\text{(TP + FP)(TP + FN)(TN + FP)(TN + FN)}}}$$

**Area under Receiver Operating Characteristic (AUROC) score:** The Receiver Operating Characteristic (ROC) curve illustrates the performance of binary and multiclass classification models at different threshold values. It is a plot of the True Positive Rate (TPR) against the False Positive Rate (FPR) for all possible threshold values. The area under the ROC curve (AuROC) indicates the model's performance, with scores ranging from 0 to 1. Higher AuROC values, closer to 1, signify better model performance.

#### 2 Supplementary Figures

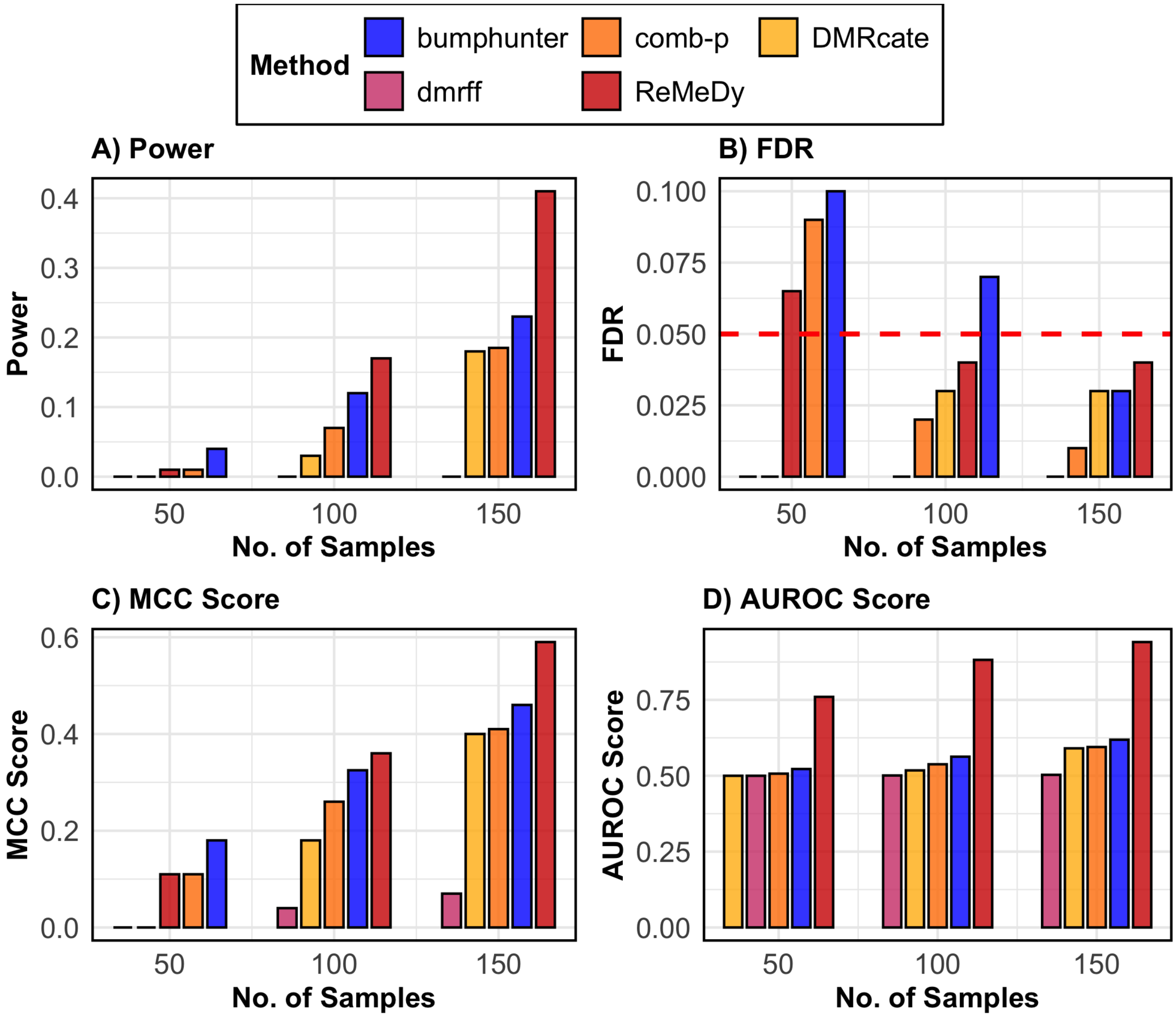

**SFigure 1: Performance of ReMeDy and competing methods in the DMR only scenario**. This panel plot compares ReMeDy with other competing methods in DMR only scenario across three sample sizes (50, 100, and 150) with mean effect of 0.4 and equal group proportions. Panel A shows statistical power, Panel B shows FDR, Panel C shows the MCC score and Panel D shows AUROC scores. Each bar reflects median of 100 simulation runs. The dashed red line in Panel B marks the nominal FDR threshold of 0.05. Higher values in Panels A, C, and D indicate better performance. Methods without a visible bar achieved value of zero.

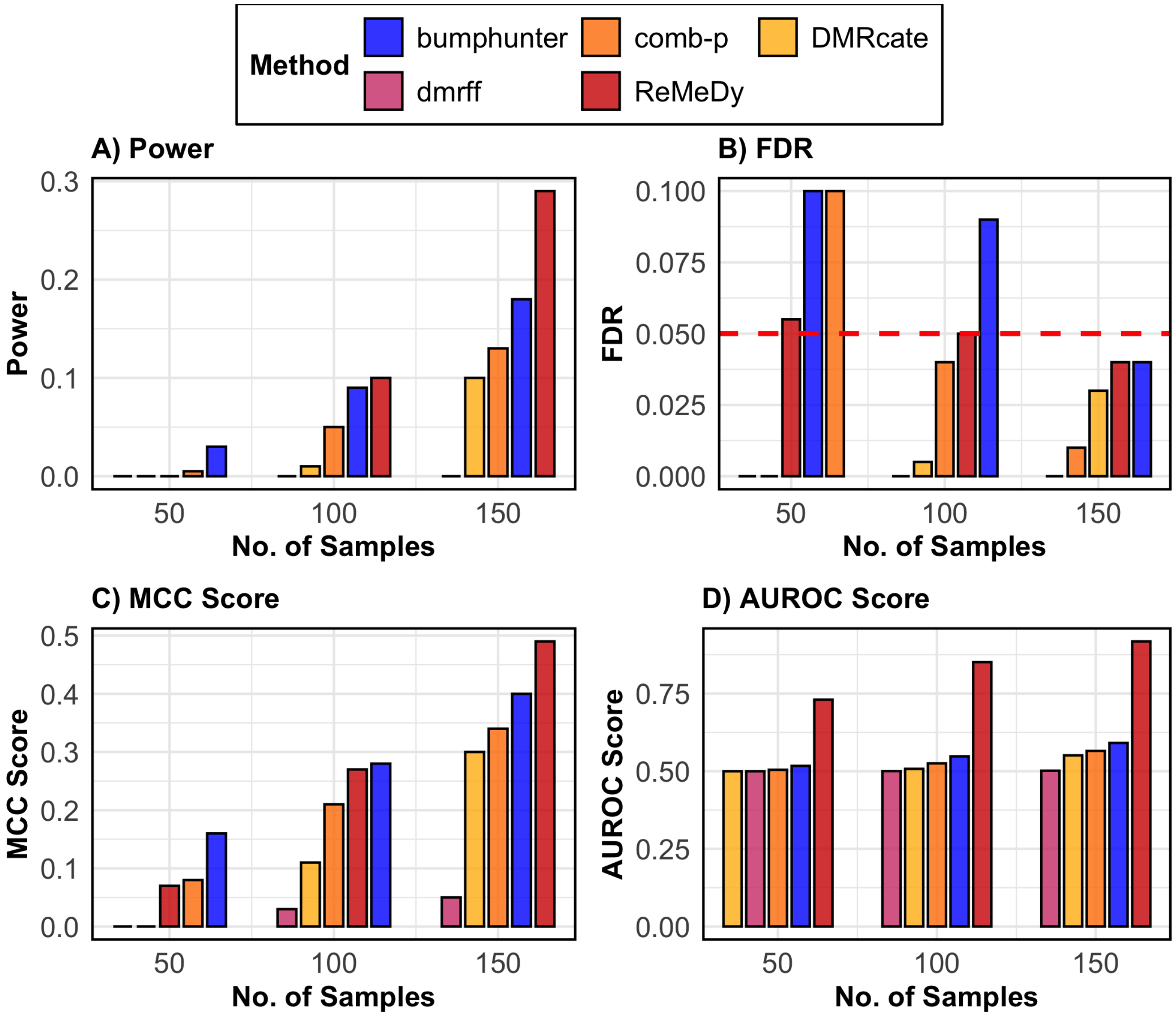

**SFigure 2: Performance of ReMeDy and competing methods in the DMR only scenario**. This panel plot compares ReMeDy with other competing methods in DMR only scenario across three sample sizes (50, 100, and 150) with mean effect of 0.4 and unequal group proportions. Panel A shows statistical power, Panel B shows FDR, Panel C shows the MCC score and Panel D shows AUROC scores. Each bar reflects median of 100 simulation runs. The dashed red line in Panel B marks the nominal FDR threshold of 0.05. Higher values in Panels A, C, and D indicate better performance. Methods without a visible bar achieved value of zero.

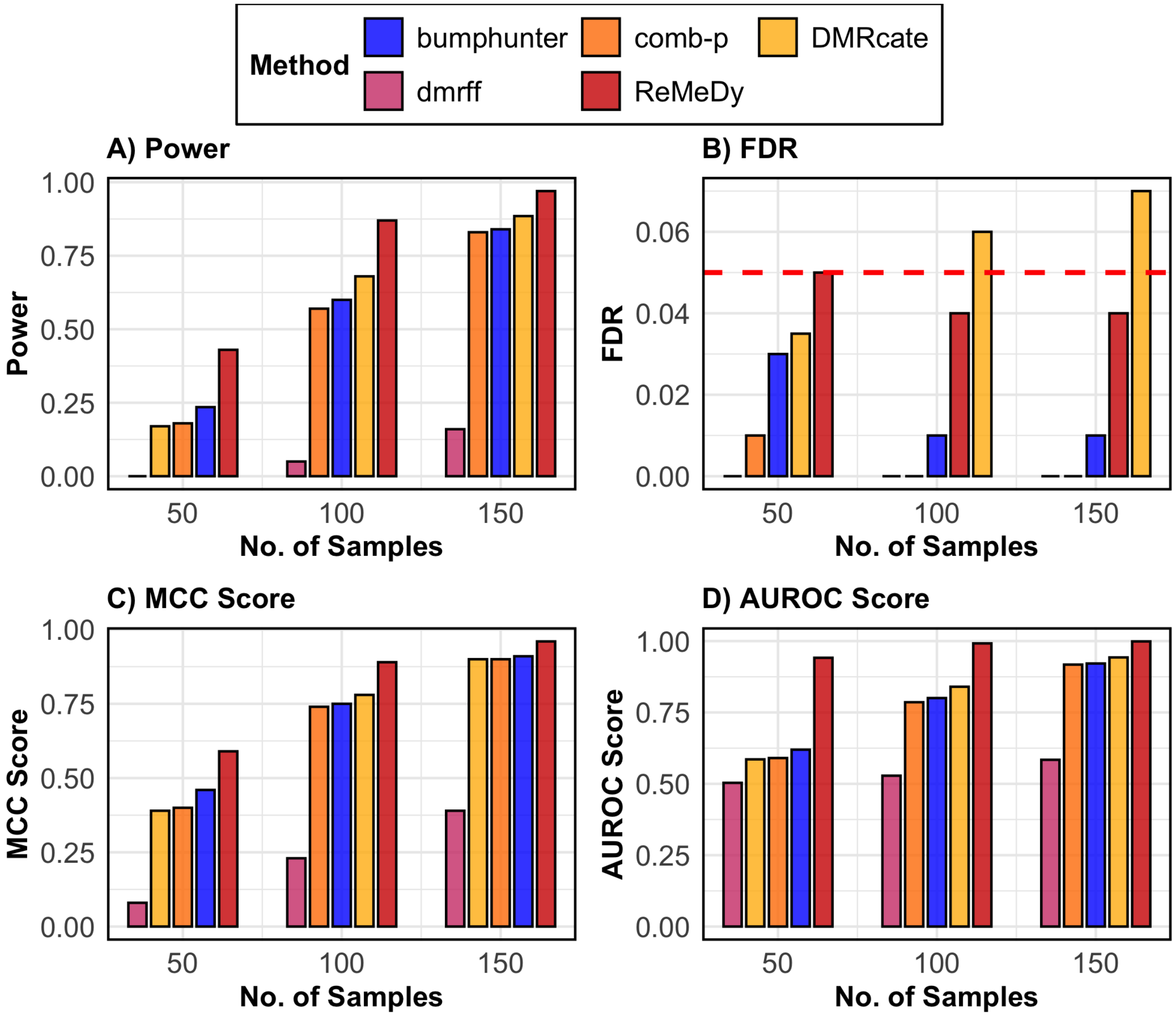

SFigure 3: **Performance of ReMeDy and competing methods in the DMR only scenario**. This panel plot compares ReMeDy with other competing methods in DMR only scenario across three sample sizes (50, 100, and 150) with mean effect of 0.7 and equal group proportions. Panel A shows statistical power, Panel B shows FDR, Panel C shows the MCC score and Panel D shows AUROC scores. Each bar reflects median of 100 simulation runs. The dashed red line in Panel B marks the nominal FDR threshold of 0.05. Higher values in Panels A, C, and D indicate better performance. Methods without a visible bar achieved value of zero.

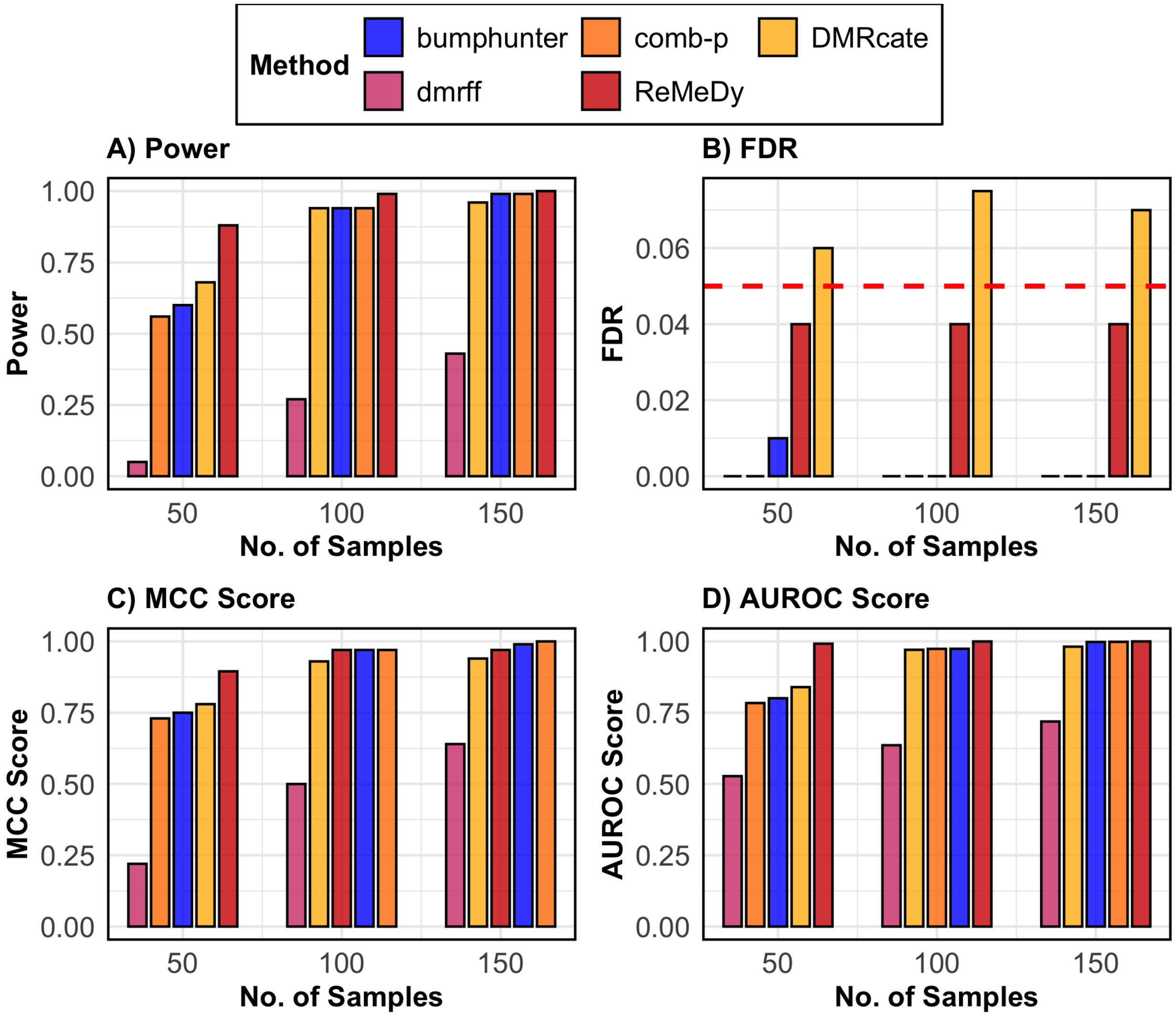

**SFigure 4: Performance of ReMeDy and competing methods in the DMR only scenario**. This panel plot compares ReMeDy with other competing methods in DMR only scenario across three sample sizes (50, 100, and 150) with mean effect of 1 and equal group proportions. Panel A shows statistical power, Panel B shows FDR, Panel C shows the MCC score and Panel D shows AUROC scores. Each bar reflects median of 100 simulation runs. The dashed red line in Panel B marks the nominal FDR threshold of 0.05. Higher values in Panels A, C, and D indicate better performance. Methods without a visible bar achieved value of zero.

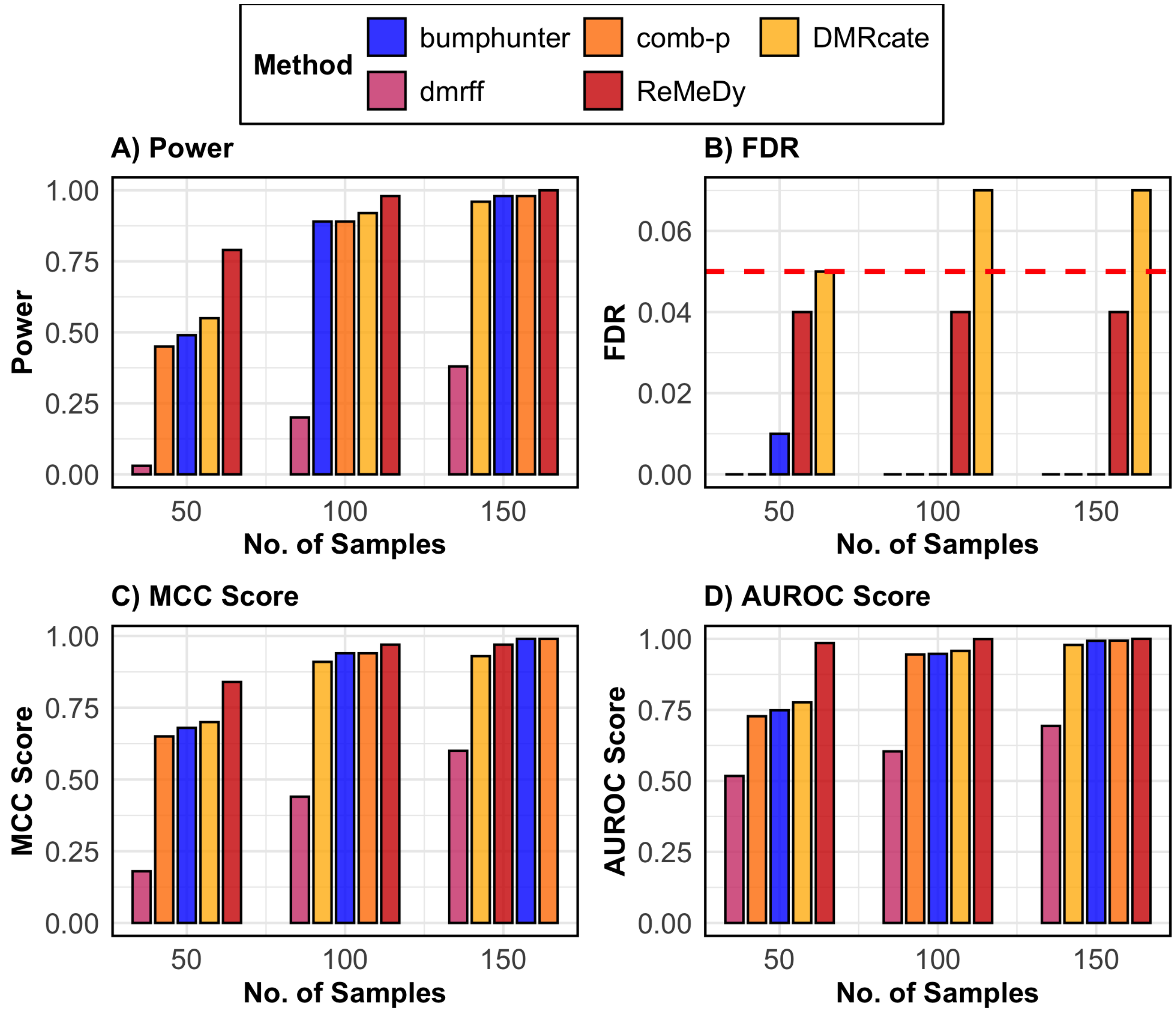

**SFigure 5: Performance of ReMeDy and competing methods in the DMR only scenario**. This panel plot compares ReMeDy with other competing methods in DMR only scenario across three sample sizes (50, 100, and 150) with mean effect of 1 and unequal group proportions. Panel A shows statistical power, Panel B shows FDR, Panel C shows the MCC score and Panel D shows AUROC scores. Each bar reflects median of 100 simulation runs. The dashed red line in Panel B marks the nominal FDR threshold of 0.05. Higher values in Panels A, C, and D indicate better performance. Methods without a visible bar achieved value of zero.

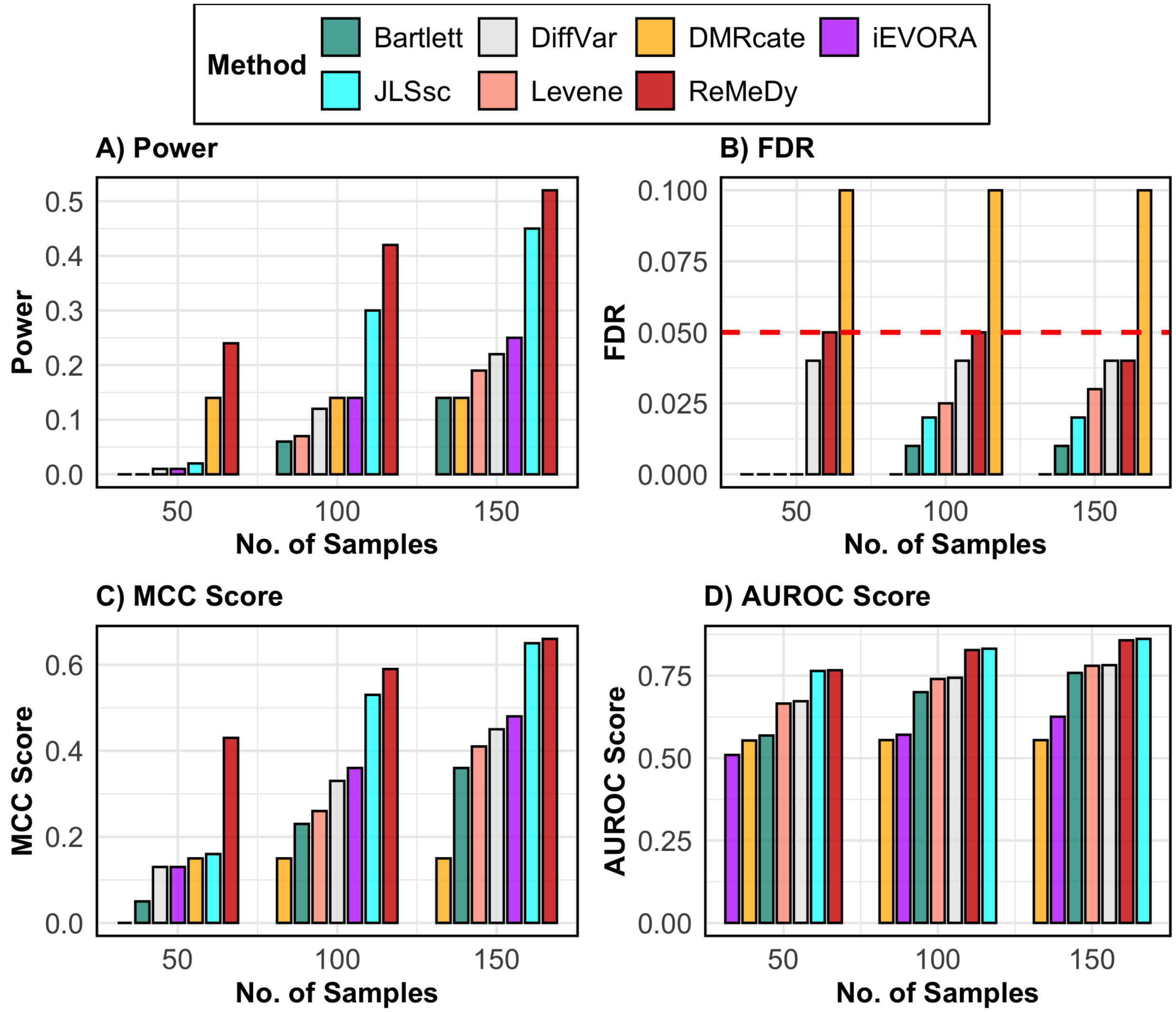

**SFigure 6: Performance of ReMeDy and competing methods in the VMR only scenario**. This panel plot compares ReMeDy with other competing methods in VMR only scenario across three sample sizes (50, 100, and 150) with variance effect of 1.5 and equal group proportions. Panel A shows statistical power, Panel B shows FDR, Panel C shows the MCC score and Panel D shows AUROC scores. Each bar reflects median of 100 simulation runs. The dashed red line in Panel B marks the nominal FDR threshold of 0.05. Higher values in Panels A, C, and D indicate better performance. Methods without a visible bar achieved value of zero.

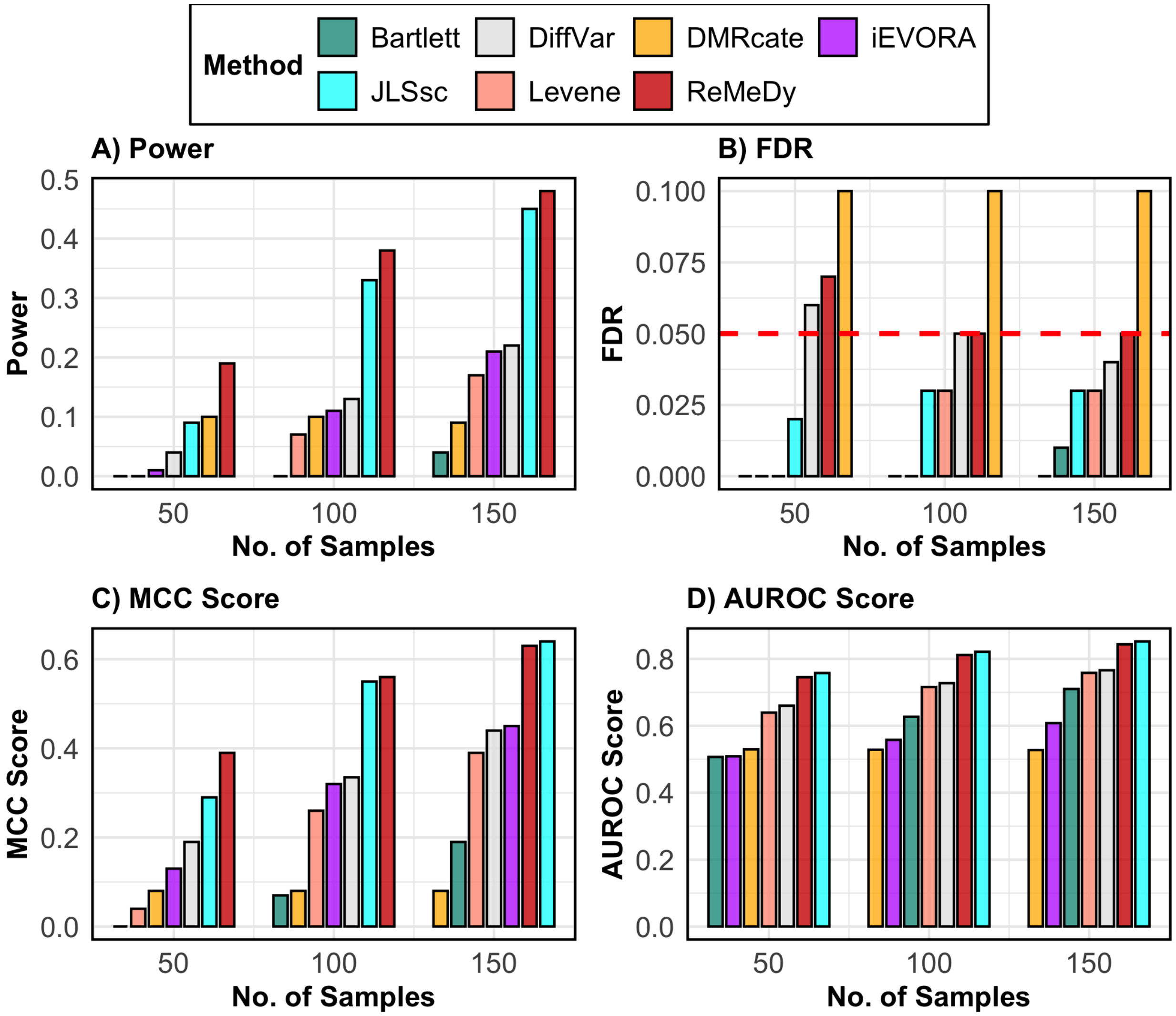

**SFigure 7**: **Performance of ReMeDy and competing methods in the VMR only scenario**. This panel plot compares ReMeDy with other competing methods in VMR only scenario across three sample sizes (50, 100, and 150) with variance effect of 1.5 and unequal group proportions. Panel A shows statistical power, Panel B shows FDR, Panel C shows the MCC score and Panel D shows AUROC scores. Each bar reflects median of 100 simulation runs. The dashed red line in Panel B marks the nominal FDR threshold of 0.05. Higher values in Panels A, C, and D indicate better performance. Methods without a visible bar achieved value of zero.

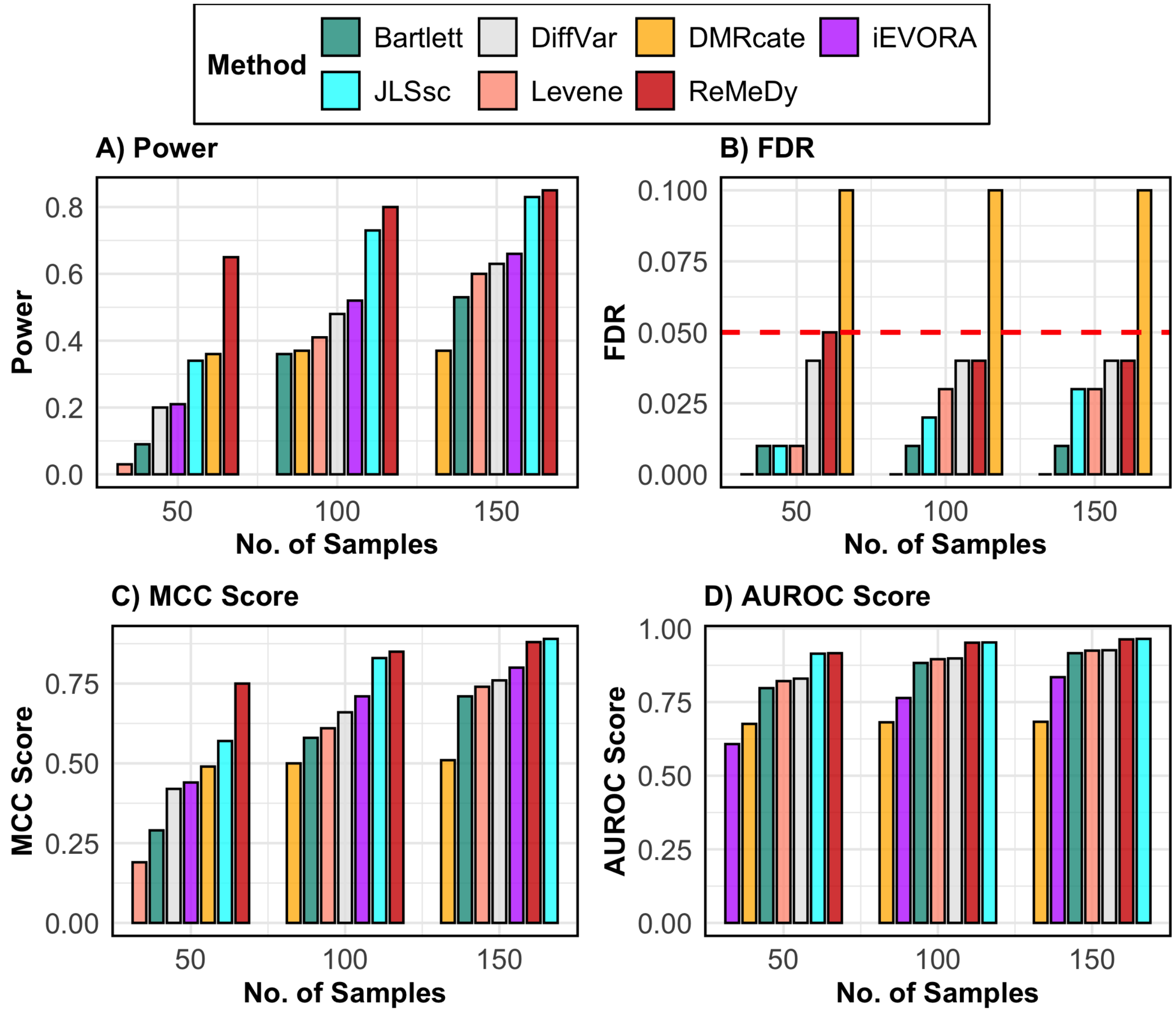

**SFigure 8**: **Performance of ReMeDy and competing methods in the VMR only scenario**. This panel plot compares ReMeDy with other competing methods in VMR only scenario across three sample sizes (50, 100, and 150) with variance effect of 2.5 and equal group proportions. Panel A shows statistical power, Panel B shows FDR, Panel C shows the MCC score and Panel D shows AUROC scores. Each bar reflects median of 100 simulation runs. The dashed red line in Panel B marks the nominal FDR threshold of 0.05. Higher values in Panels A, C, and D indicate better performance. Methods without a visible bar achieved value of zero.

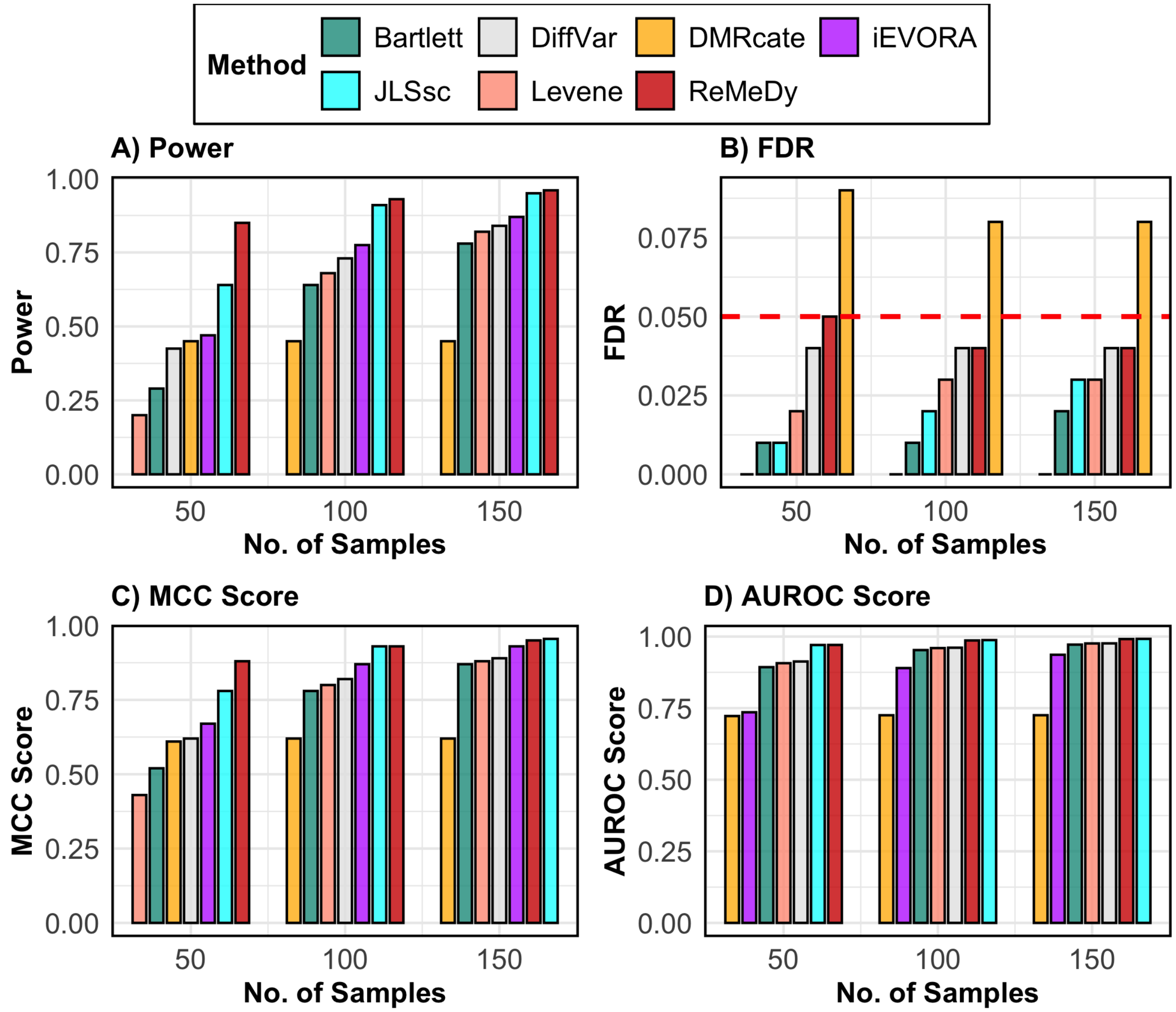

**SFigure 9: Performance of ReMeDy and competing methods in the VMR only scenario**. This panel plot compares ReMeDy with other competing methods in VMR only scenario across three sample sizes (50, 100, and 150) with variance effect of 3.5 and equal group proportions. Panel A shows statistical power, Panel B shows FDR, Panel C shows the MCC score and Panel D shows AUROC scores. Each bar reflects median of 100 simulation runs. The dashed red line in Panel B marks the nominal FDR threshold of 0.05. Higher values in Panels A, C, and D indicate better performance. Methods without a visible bar achieved value of zero.

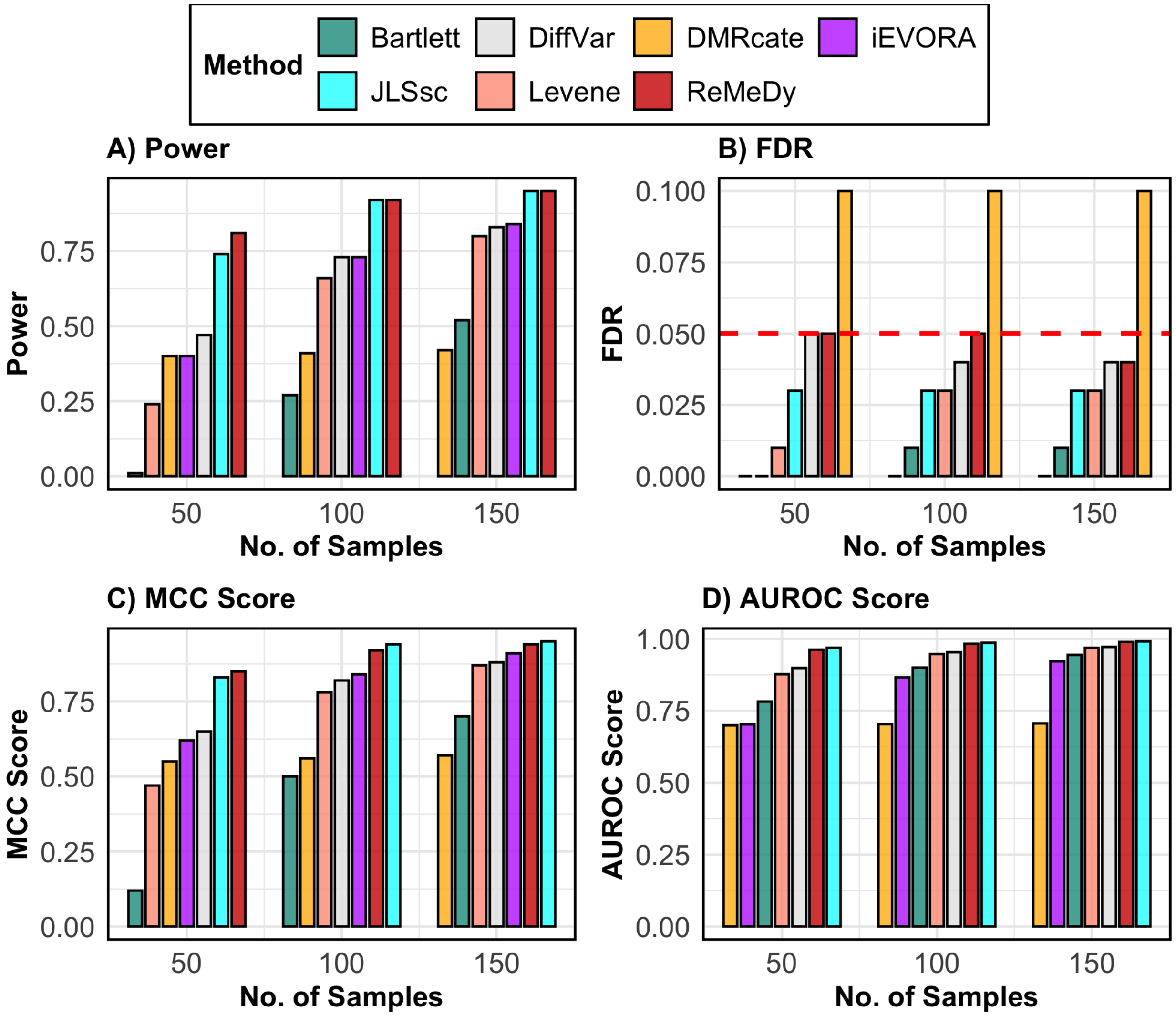

**SFigure** 10: **Performance of ReMeDy and competing methods in the VMR only scenario**. This panel plot compares ReMeDy with other competing methods in VMR only scenario across three sample sizes (50, 100, and 150) with variance effect of 3.5 and unequal group proportions. Panel A shows statistical power, Panel B shows FDR, Panel C shows the MCC score and Panel D shows AUROC scores. Each bar reflects median of 100 simulation runs. The dashed red line in Panel B marks the nominal FDR threshold of 0.05. Higher values in Panels A, C, and D indicate better performance. Methods without a visible bar achieved value of zero.

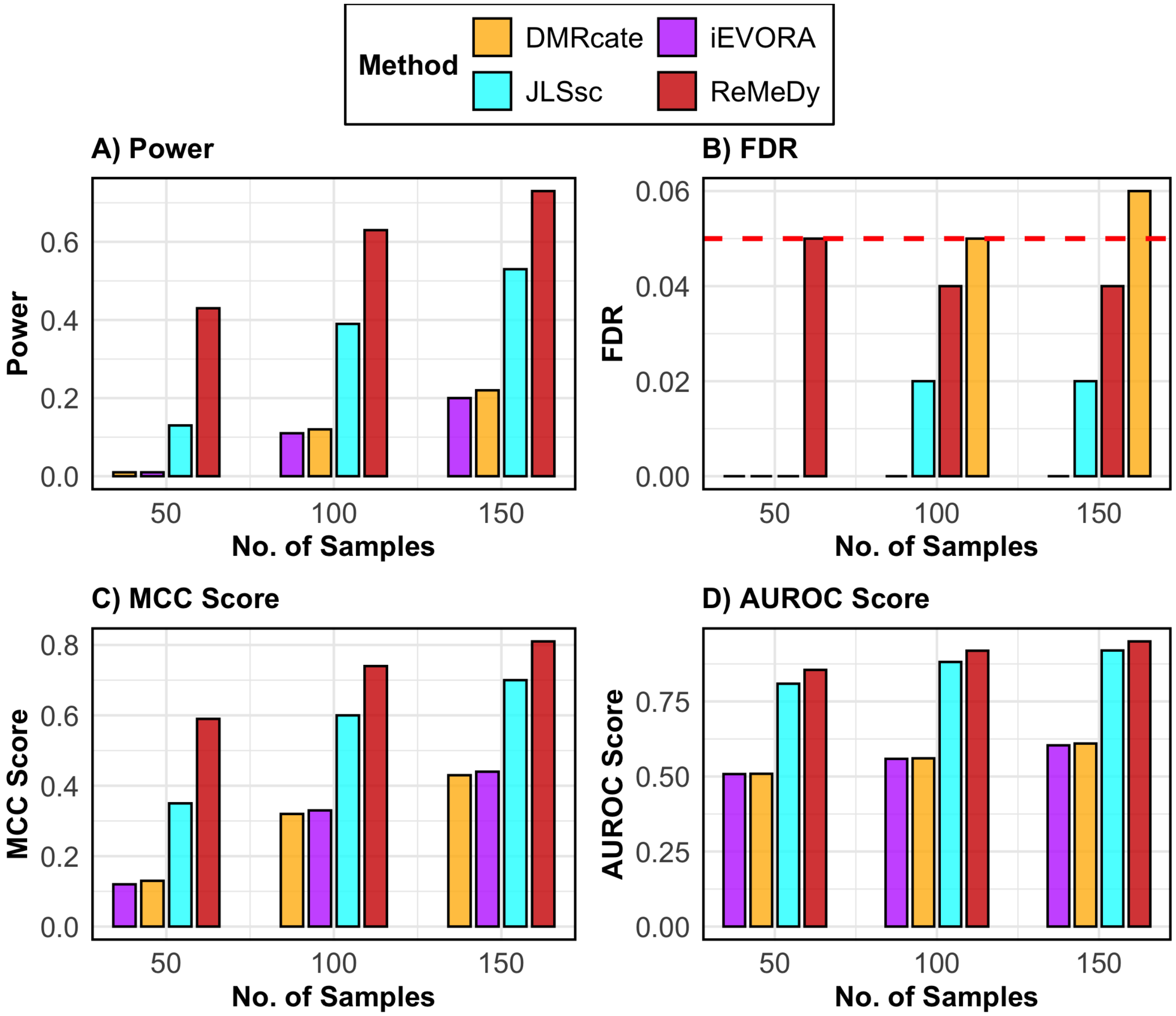

**SFigure** 11: **Performance of ReMeDy and competing methods in the DVMR scenario**. This panel plot compares ReMeDy with other competing methods in DVMR scenario across three sample sizes (50, 100, and 150) with mean effect of 0.4, variance effect of 1.5 and equal group proportions. Panel A shows statistical power, Panel B shows FDR, Panel C shows the MCC score and Panel D shows AUROC scores. Each bar reflects median of 100 simulation runs. The dashed red line in Panel B marks the nominal FDR threshold of 0.05. Higher values in Panels A, C, and D indicate better performance. Methods without a visible bar achieved value of zero.

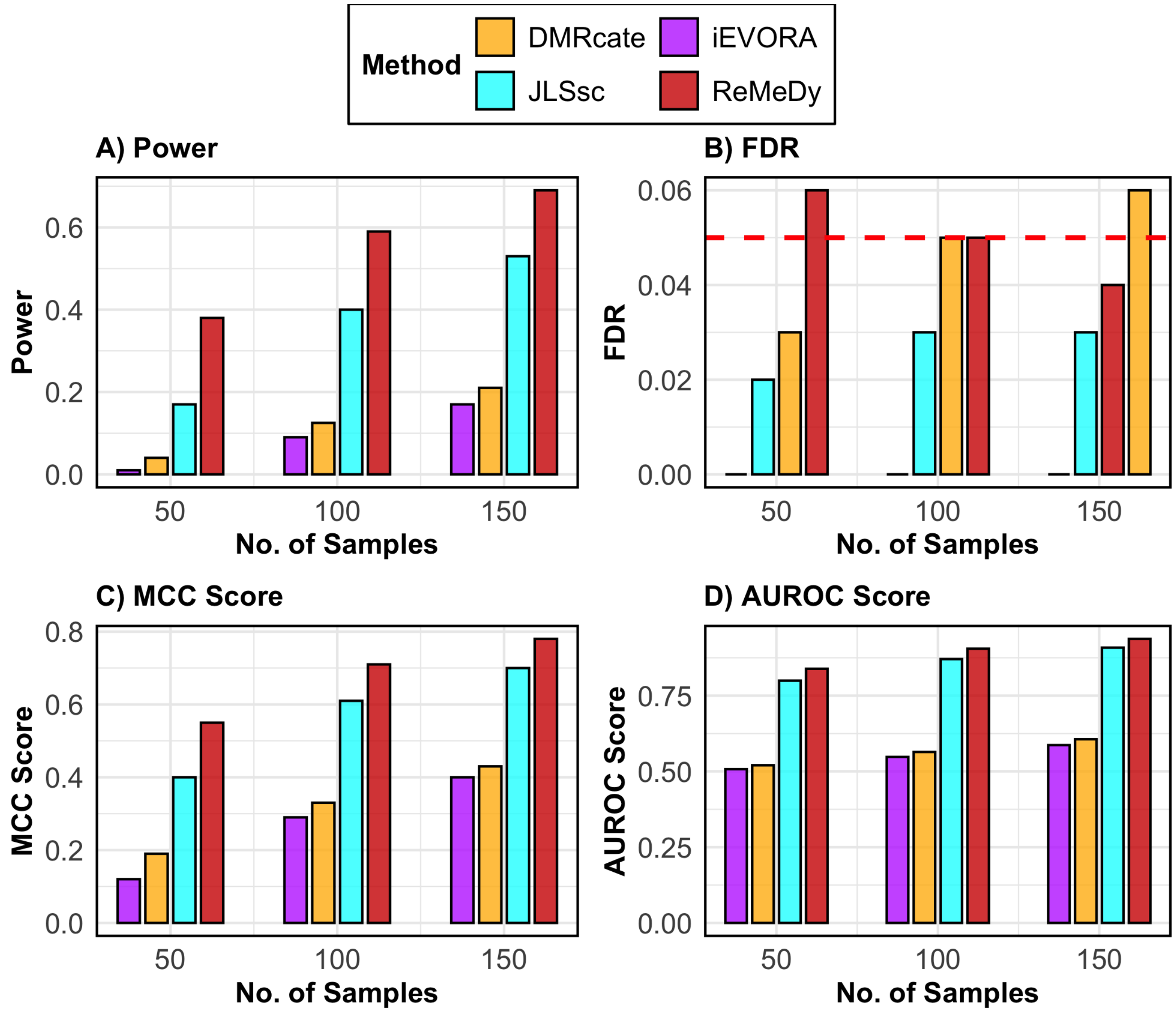

**SFigure** 12: **Performance of ReMeDy and competing methods in the DVMR scenario**. This panel plot compares ReMeDy with other competing methods in DVMR scenario across three sample sizes (50, 100, and 150) with mean effect of 0.4, variance effect of 1.5 and unequal group proportions. Panel A shows statistical power, Panel B shows FDR, Panel C shows the MCC score and Panel D shows AUROC scores. Each bar reflects median of 100 simulation runs. The dashed red line in Panel B marks the nominal FDR threshold of 0.05. Higher values in Panels A, C, and D indicate better performance. Methods without a visible bar achieved value of zero.

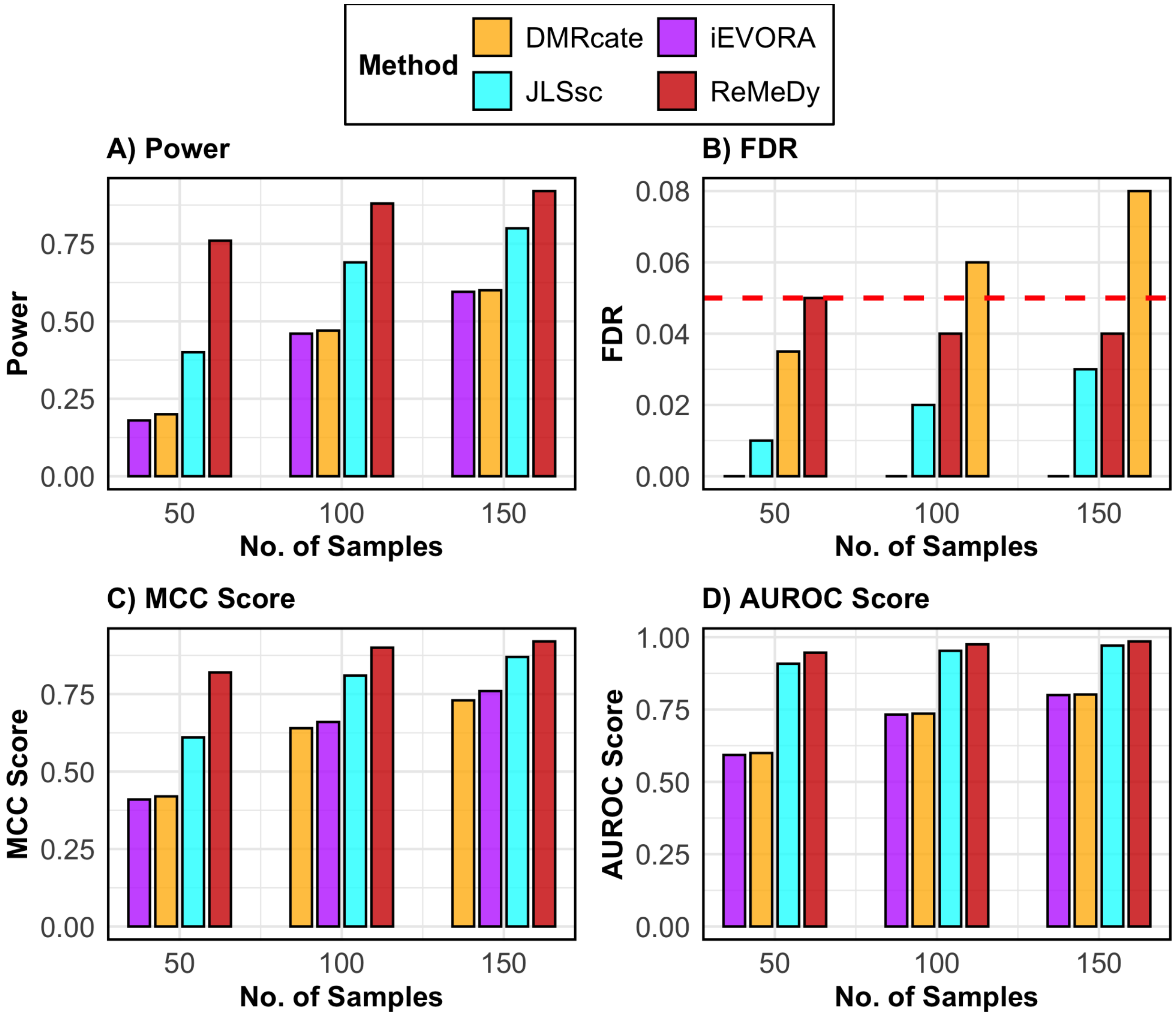

**SFigure** 13: **Performance of ReMeDy and competing methods in the DVMR scenario**. This panel plot compares ReMeDy with other competing methods in DVMR scenario across three sample sizes (50, 100, and 150) with mean effect of 0.4, variance effect of 2.5 and equal group proportions. Panel A shows statistical power, Panel B shows FDR, Panel C shows the MCC score and Panel D shows AUROC scores. Each bar reflects median of 100 simulation runs. The dashed red line in Panel B marks the nominal FDR threshold of 0.05. Higher values in Panels A, C, and D indicate better performance. Methods without a visible bar achieved value of zero.

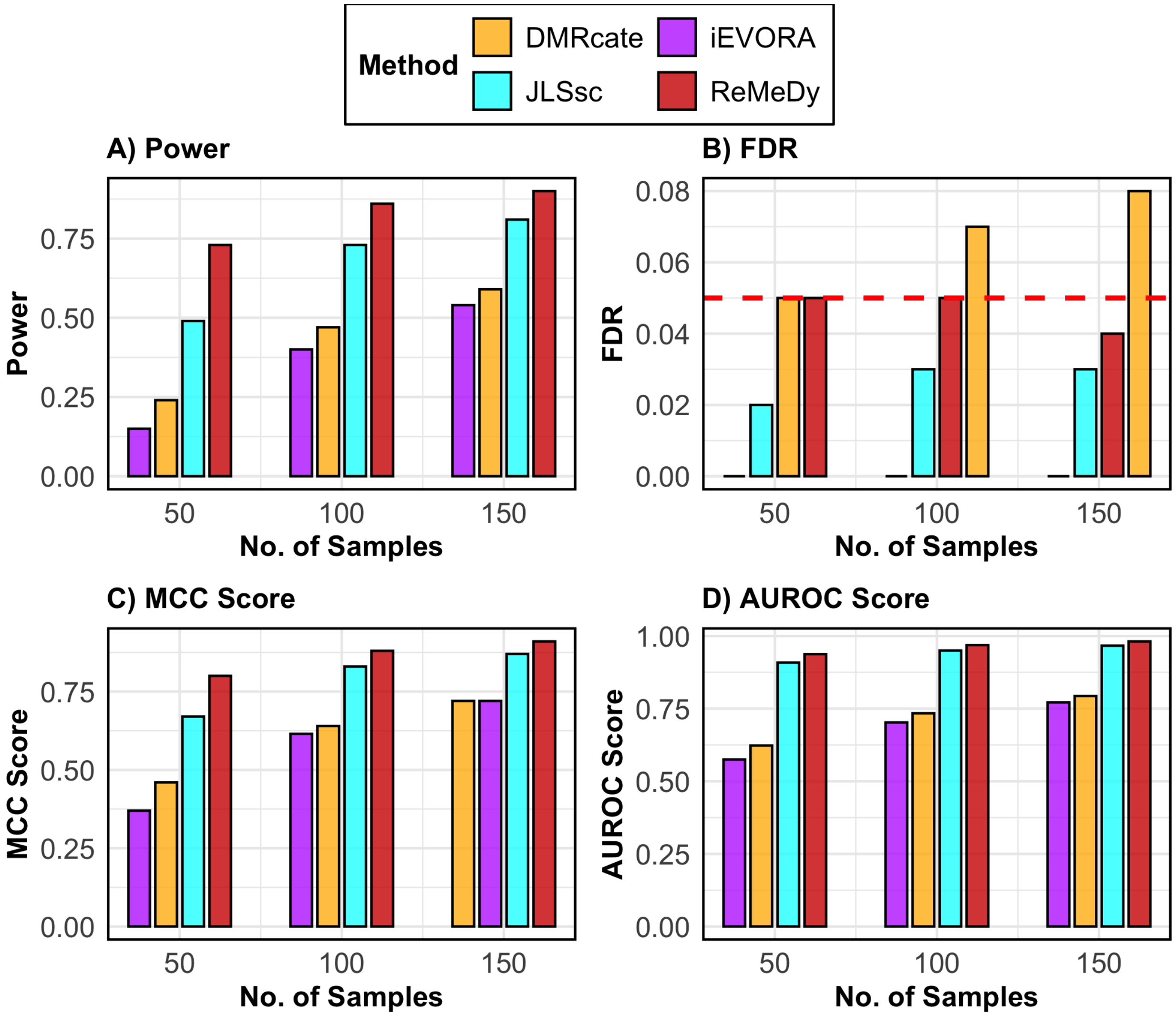

**SFigure** 14: **Performance of ReMeDy and competing methods in the DVMR scenario**. This panel plot compares ReMeDy with other competing methods in DVMR scenario across three sample sizes (50, 100, and 150) with mean effect of 0.4, variance effect of 2.5 and unequal group proportions. Panel A shows statistical power, Panel B shows FDR, Panel C shows the MCC score and Panel D shows AUROC scores. Each bar reflects median of 100 simulation runs. The dashed red line in Panel B marks the nominal FDR threshold of 0.05. Higher values in Panels A, C, and D indicate better performance. Methods without a visible bar achieved value of zero.

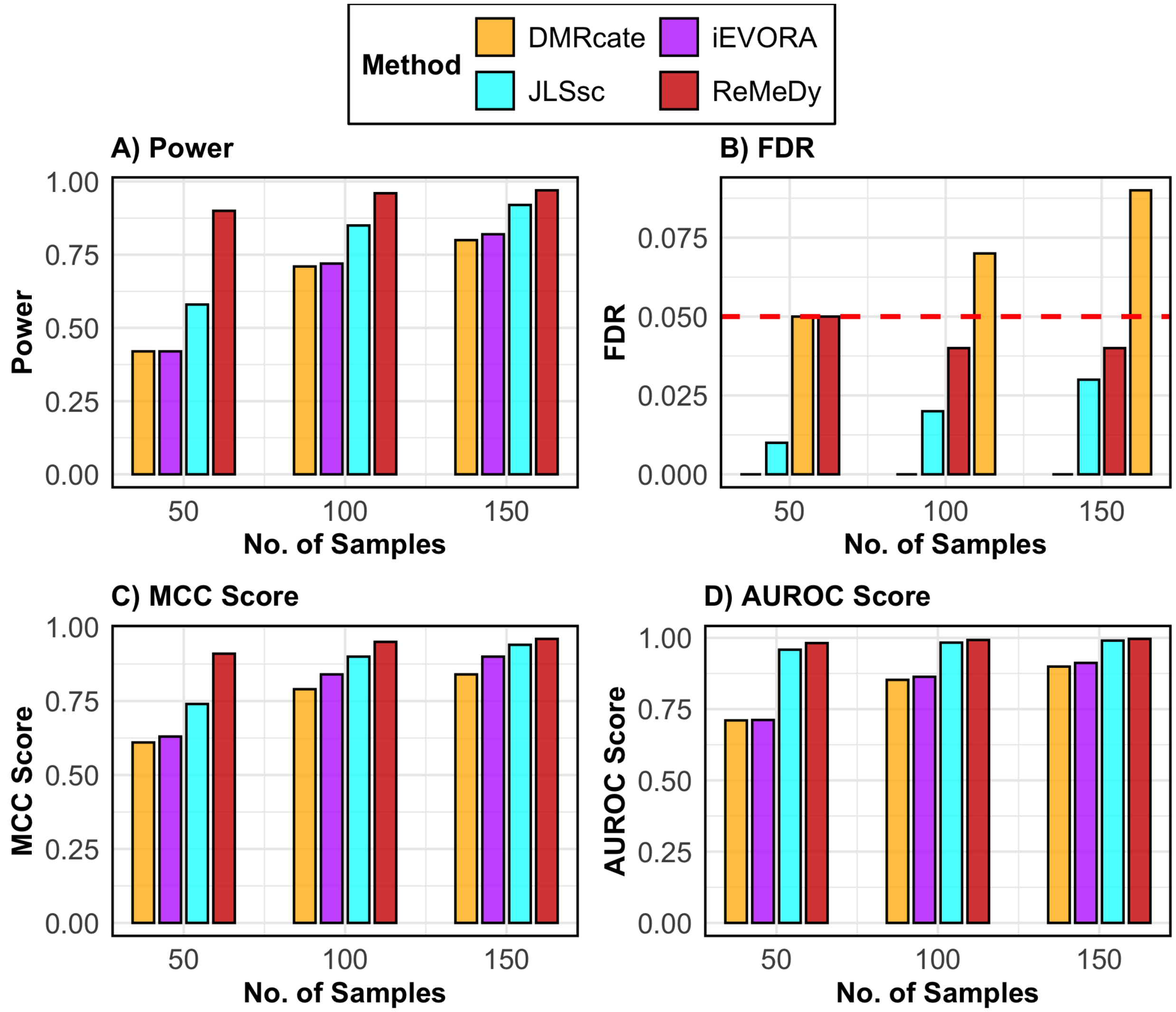

**SFigure** 15: **Performance of ReMeDy and competing methods in the DVMR scenario**. This panel plot compares ReMeDy with other competing methods in DVMR scenario across three sample sizes (50, 100, and 150) with mean effect of 0.4, variance effect of 3.5 and equal group proportions. Panel A shows statistical power, Panel B shows FDR, Panel C shows the MCC score and Panel D shows AUROC scores. Each bar reflects median of 100 simulation runs. The dashed red line in Panel B marks the nominal FDR threshold of 0.05. Higher values in Panels A, C, and D indicate better performance. Methods without a visible bar achieved value of zero.

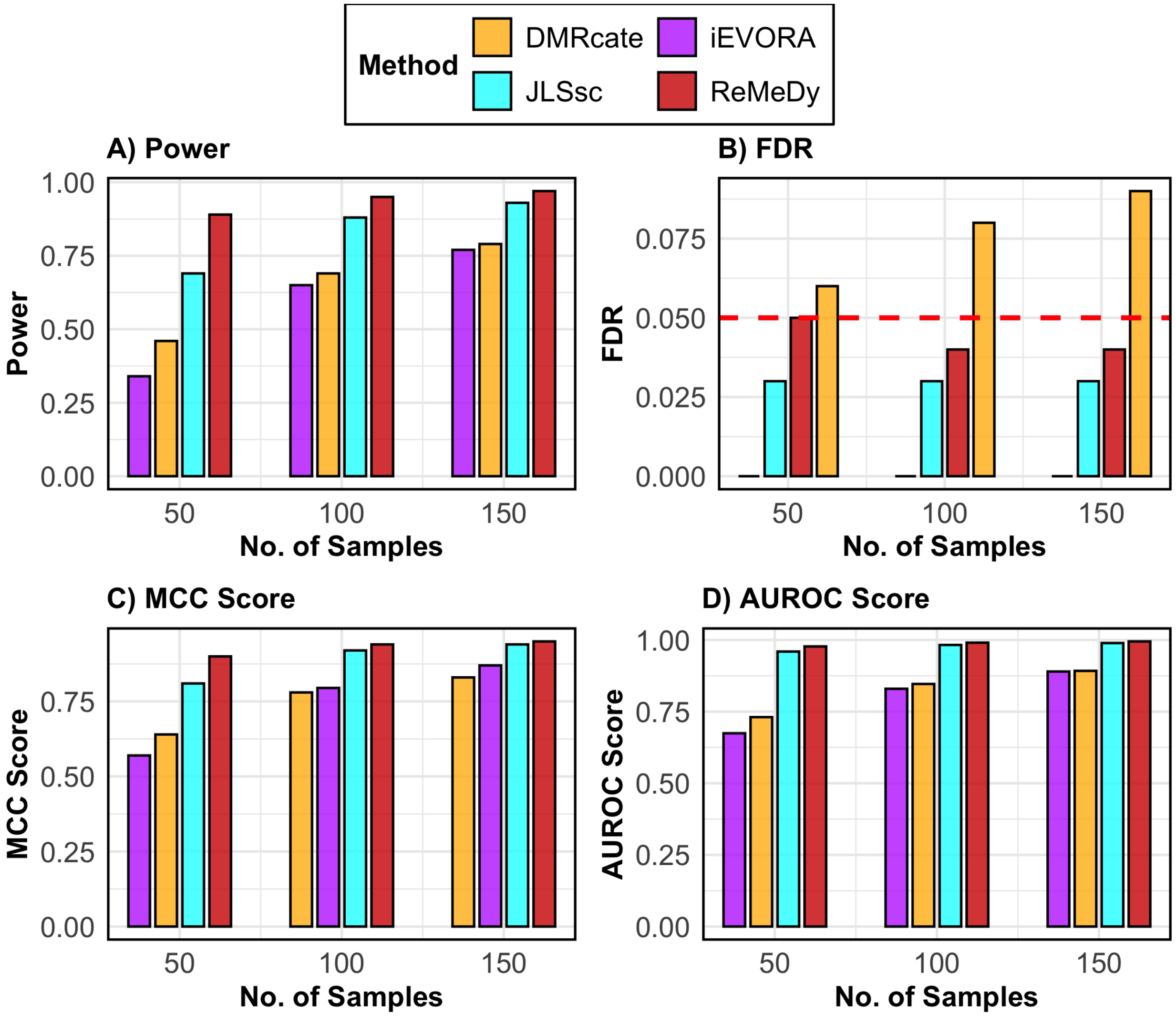

**SFigure** 16: **Performance of ReMeDy and competing methods in the DVMR scenario**. This panel plot compares ReMeDy with other competing methods in DVMR scenario across three sample sizes (50, 100, and 150) with mean effect of 0.4, variance effect of 3.5 and unequal group proportions. Panel A shows statistical power, Panel B shows FDR, Panel C shows the MCC score and Panel D shows AUROC scores. Each bar reflects median of 100 simulation runs. The dashed red line in Panel B marks the nominal FDR threshold of 0.05. Higher values in Panels A, C, and D indicate better performance. Methods without a visible bar achieved value of zero.

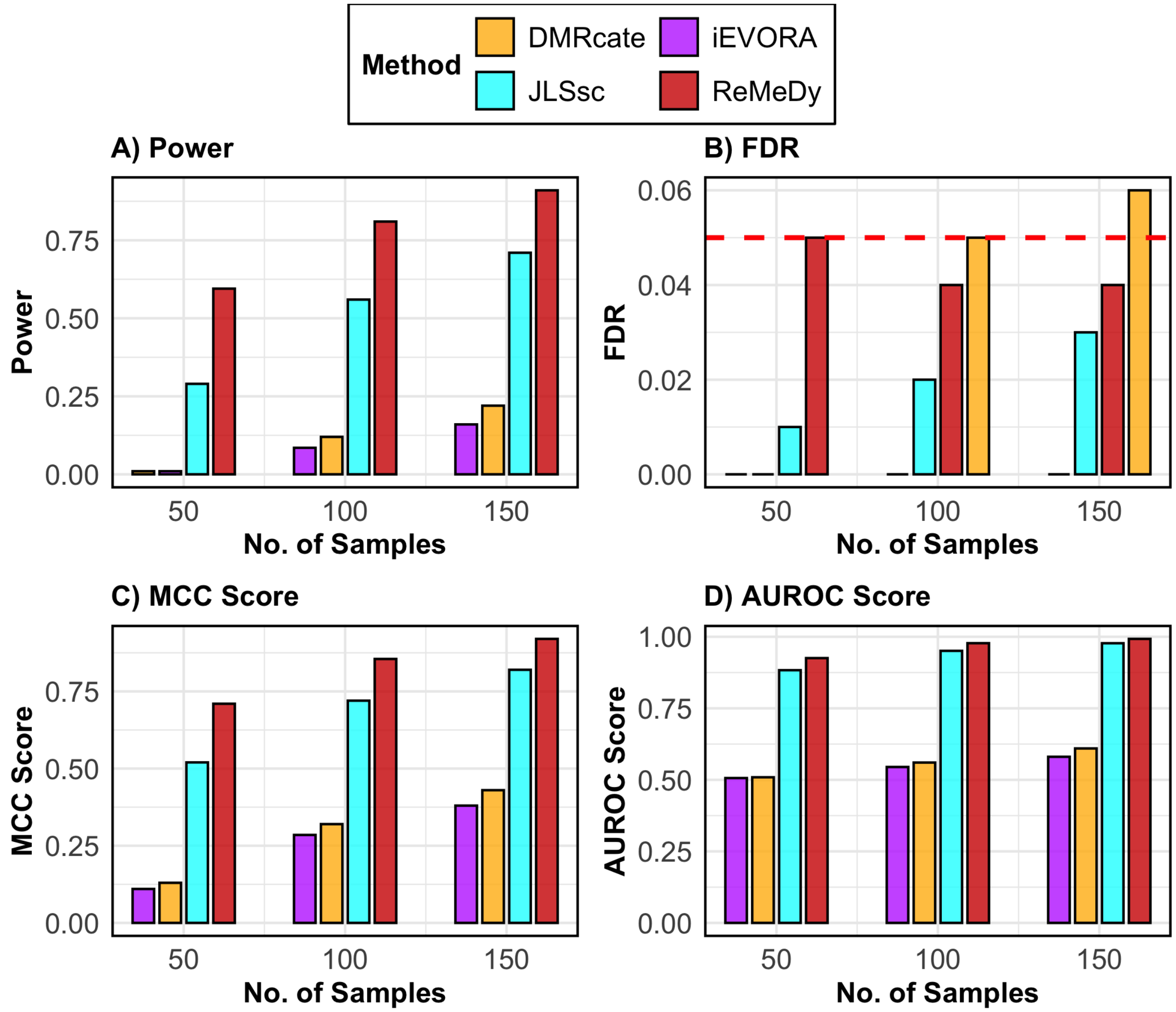

**SFigure** 17: **Performance of ReMeDy and competing methods in the DVMR scenario**. This panel plot compares ReMeDy with other competing methods in DVMR scenario across three sample sizes (50, 100, and 150) with mean effect of 0.7, variance effect of 1.5 and equal group proportions. Panel A shows statistical power, Panel B shows FDR, Panel C shows the MCC score and Panel D shows AUROC scores. Each bar reflects median of 100 simulation runs. The dashed red line in Panel B marks the nominal FDR threshold of 0.05. Higher values in Panels A, C, and D indicate better performance. Methods without a visible bar achieved value of zero.

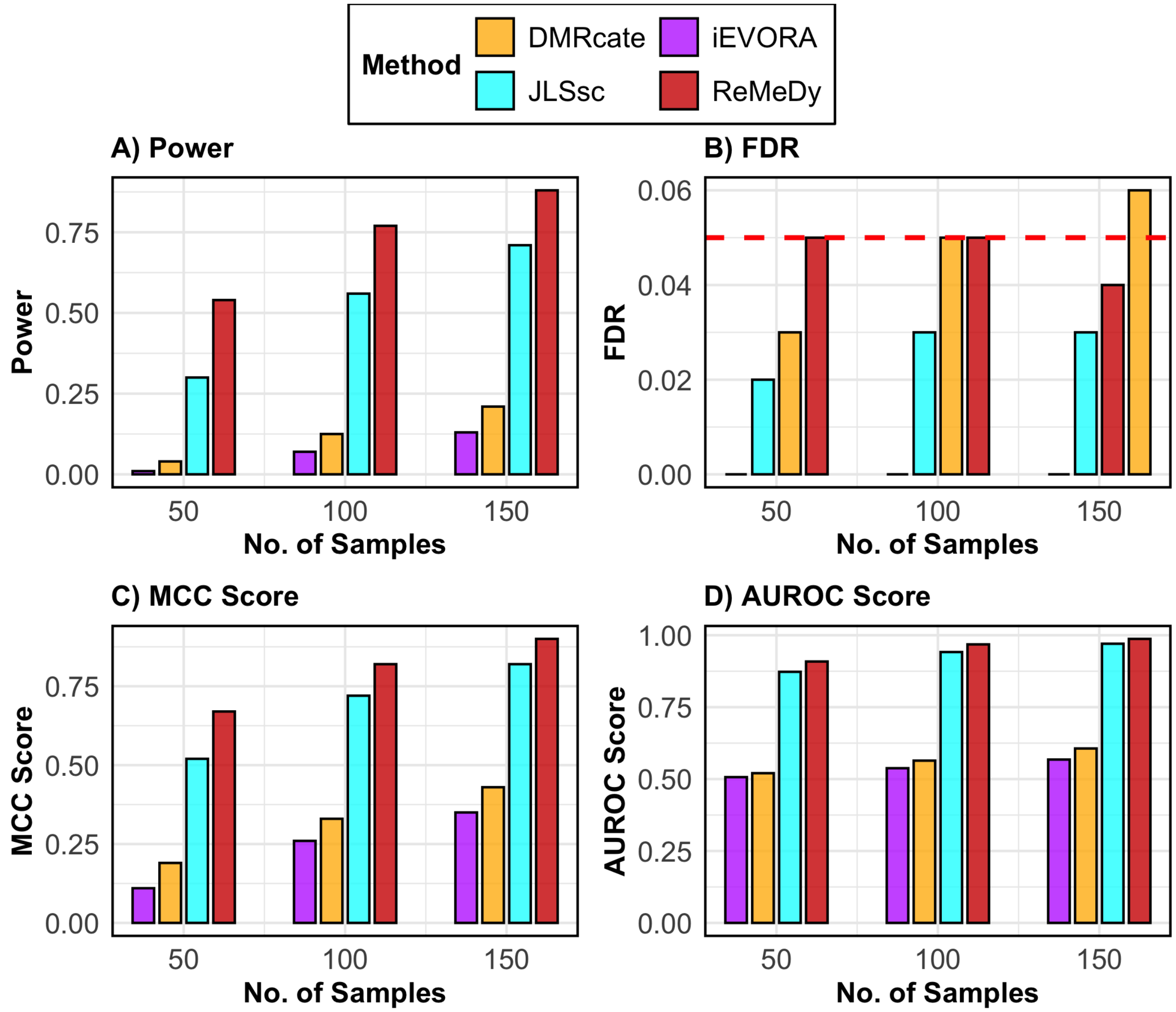

**SFigure** 18: **Performance of ReMeDy and competing methods in the DVMR scenario**. This panel plot compares ReMeDy with other competing methods in DVMR scenario across three sample sizes (50, 100, and 150) with mean effect of 0.7, variance effect of 1.5 and unequal group proportions. Panel A shows statistical power, Panel B shows FDR, Panel C shows the MCC score and Panel D shows AUROC scores. Each bar reflects median of 100 simulation runs. The dashed red line in Panel B marks the nominal FDR threshold of 0.05. Higher values in Panels A, C, and D indicate better performance. Methods without a visible bar achieved value of zero.

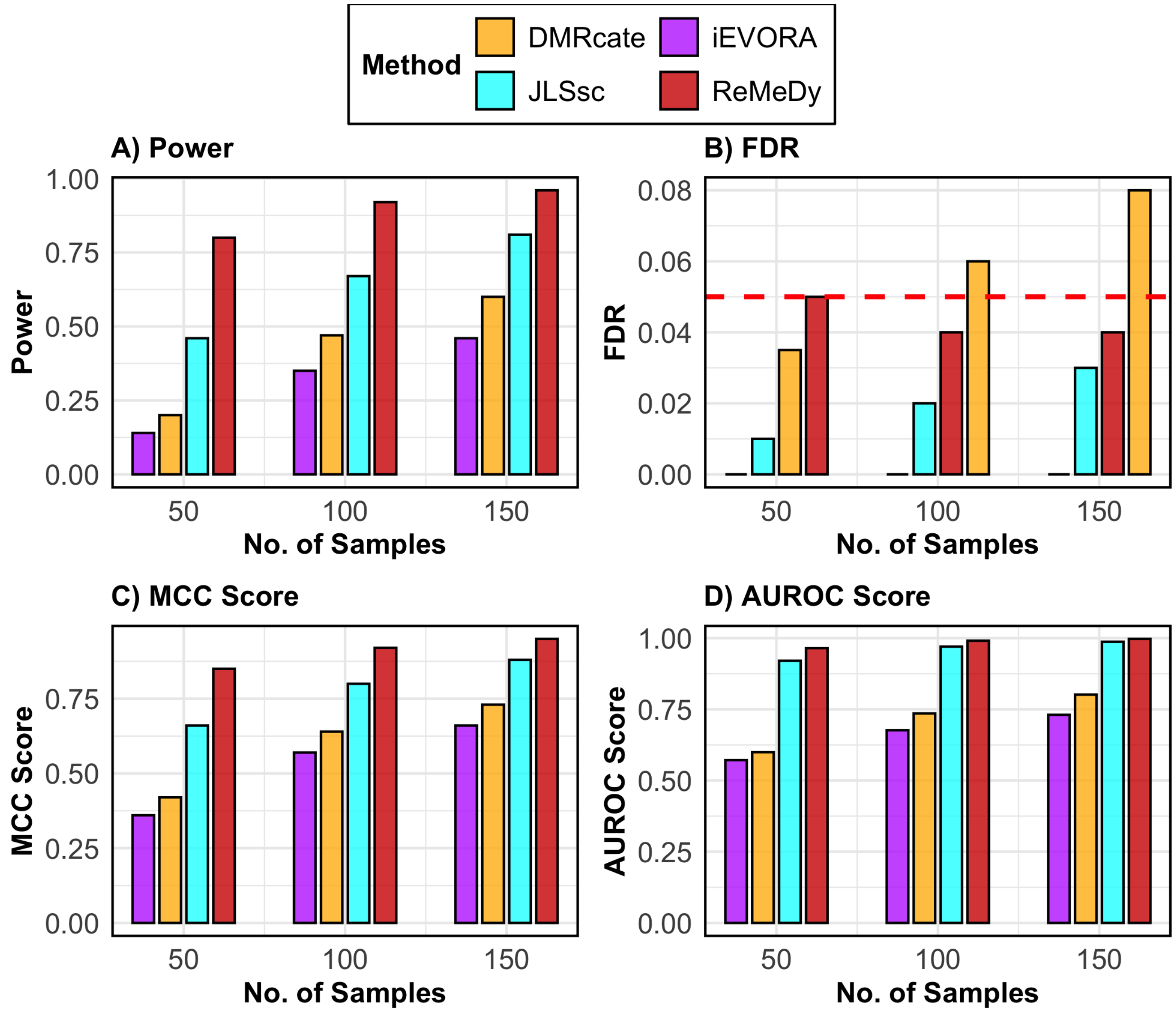

**SFigure** 19: **Performance of ReMeDy and competing methods in the DVMR scenario**. This panel plot compares ReMeDy with other competing methods in DVMR scenario across three sample sizes (50, 100, and 150) with mean effect of 0.7, variance effect of 2.5 and equal group proportions. Panel A shows statistical power, Panel B shows FDR, Panel C shows the MCC score and Panel D shows AUROC scores. Each bar reflects median of 100 simulation runs. The dashed red line in Panel B marks the nominal FDR threshold of 0.05. Higher values in Panels A, C, and D indicate better performance. Methods without a visible bar achieved value of zero.

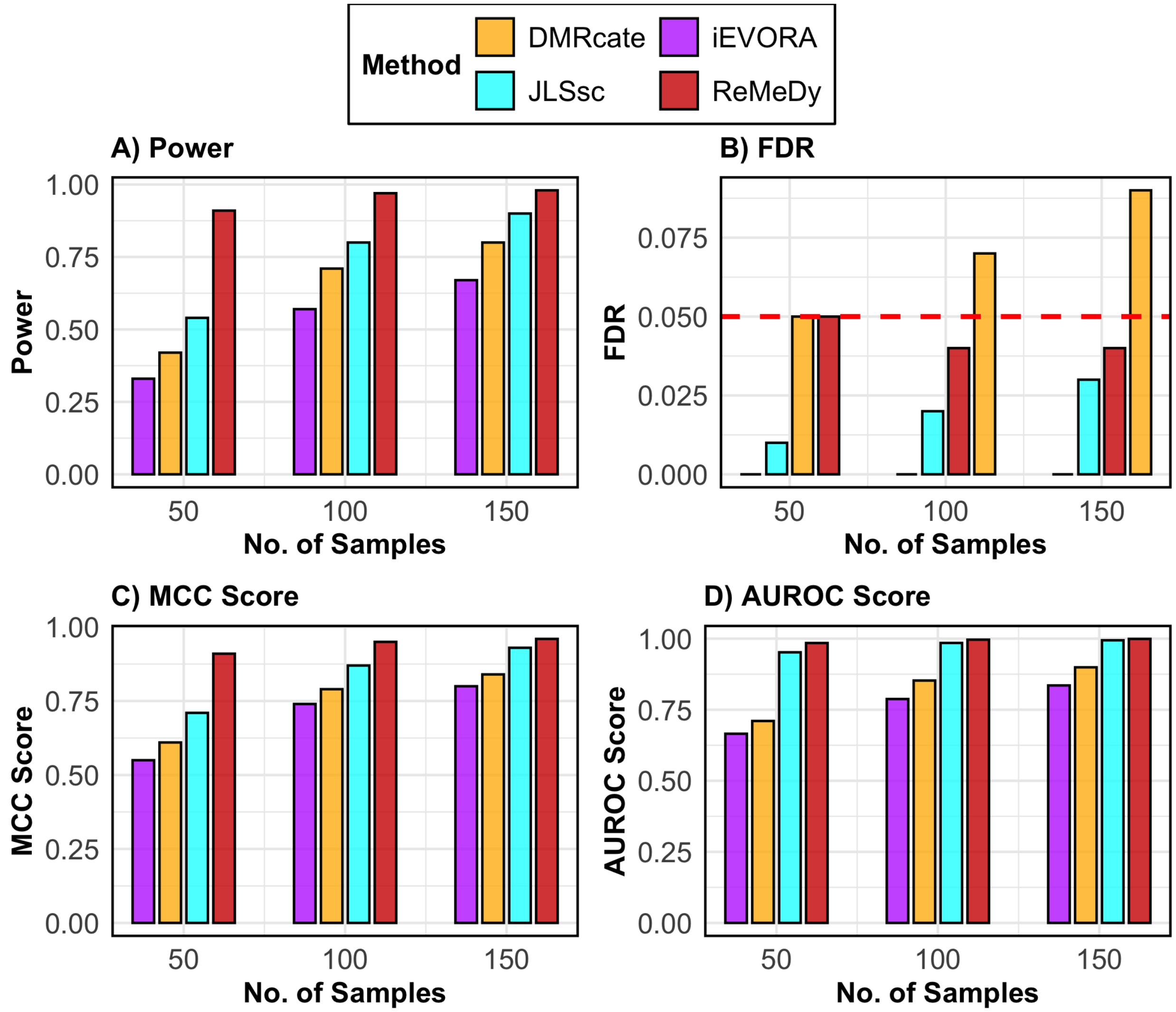

**SFigure** 20: **Performance of ReMeDy and competing methods in the DVMR scenario**. This panel plot compares ReMeDy with other competing methods in DVMR scenario across three sample sizes (50, 100, and 150) with mean effect of 0.7, variance effect of 3.5 and equal group proportions. Panel A shows statistical power, Panel B shows FDR, Panel C shows the MCC score and Panel D shows AUROC scores. Each bar reflects median of 100 simulation runs. The dashed red line in Panel B marks the nominal FDR threshold of 0.05. Higher values in Panels A, C, and D indicate better performance. Methods without a visible bar achieved value of zero.

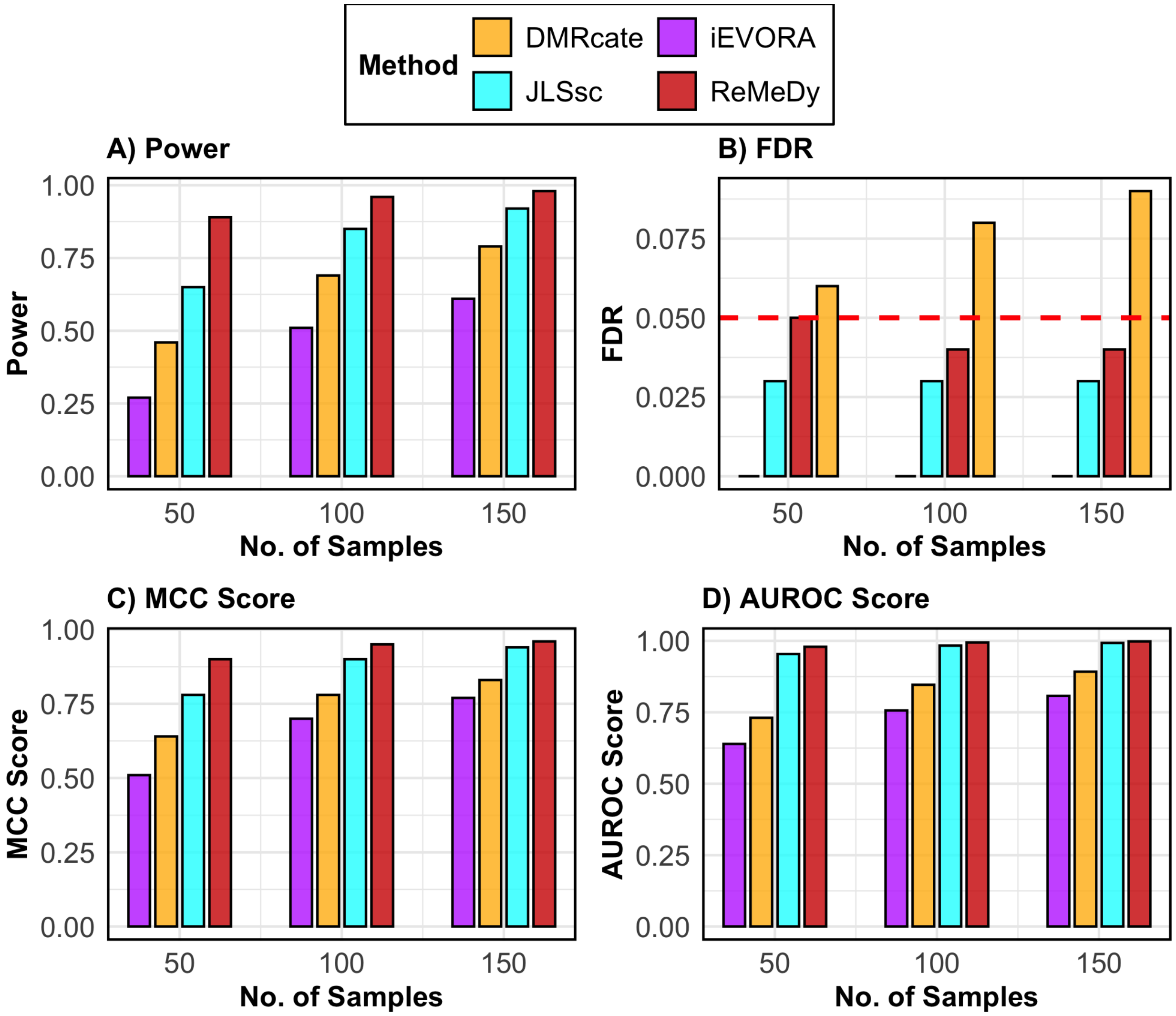

**SFigure** 21: **Performance of ReMeDy and competing methods in the DVMR scenario**. This panel plot compares ReMeDy with other competing methods in DVMR scenario across three sample sizes (50, 100, and 150) with mean effect of 0.7, variance effect of 3.5 and unequal group proportions. Panel A shows statistical power, Panel B shows FDR, Panel C shows the MCC score and Panel D shows AUROC scores. Each bar reflects median of 100 simulation runs. The dashed red line in Panel B marks the nominal FDR threshold of 0.05. Higher values in Panels A, C, and D indicate better performance. Methods without a visible bar achieved value of zero.

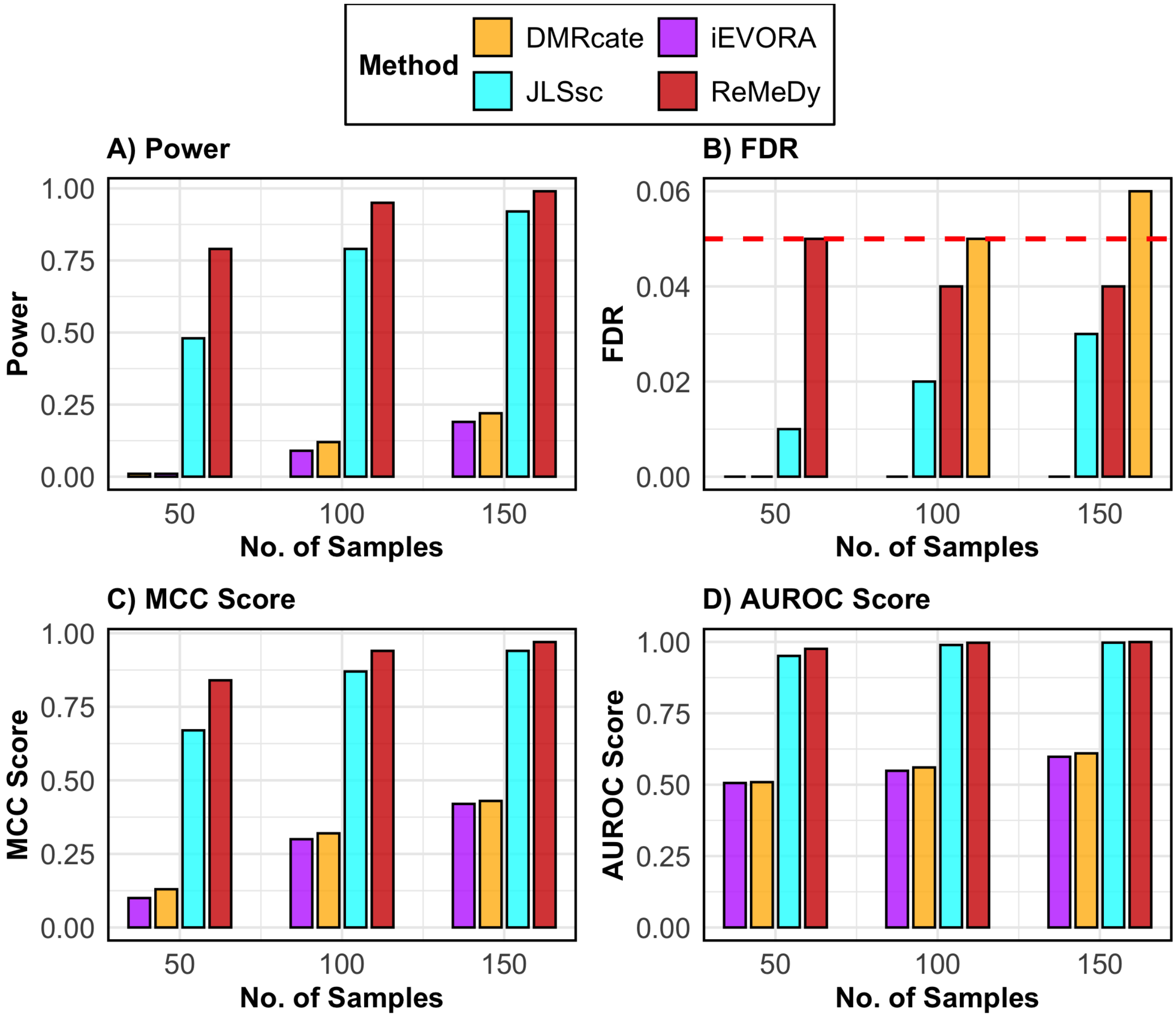

**SFigure** 22: **Performance of ReMeDy and competing methods in the DVMR scenario**. This panel plot compares ReMeDy with other competing methods in DVMR scenario across three sample sizes (50, 100, and 150) with mean effect of 1, variance effect of 1.5 and equal group proportions. Panel A shows statistical power, Panel B shows FDR, Panel C shows the MCC score and Panel D shows AUROC scores. Each bar reflects median of 100 simulation runs. The dashed red line in Panel B marks the nominal FDR threshold of 0.05. Higher values in Panels A, C, and D indicate better performance. Methods without a visible bar achieved value of zero.

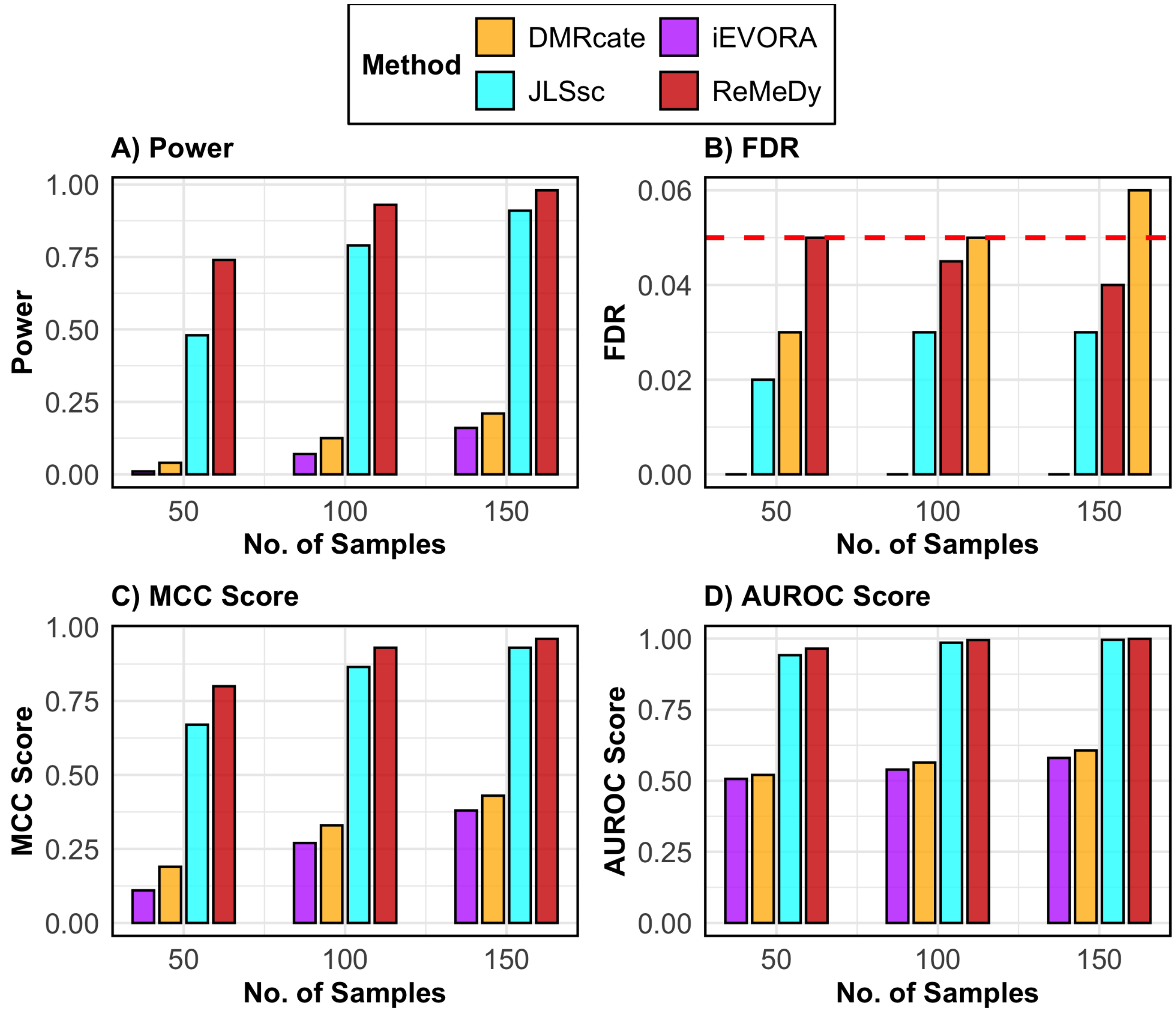

**SFigure 23: Performance of ReMeDy and competing methods in the DVMR scenario**. This panel plot compares ReMeDy with other competing methods in DVMR scenario across three sample sizes (50, 100, and 150) with mean effect of 1, variance effect of 1.5 and unequal group proportions. Panel A shows statistical power, Panel B shows FDR, Panel C shows the MCC score and Panel D shows AUROC scores. Each bar reflects median of 100 simulation runs. The dashed red line in Panel B marks the nominal FDR threshold of 0.05. Higher values in Panels A, C, and D indicate better performance. Methods without a visible bar achieved value of zero.

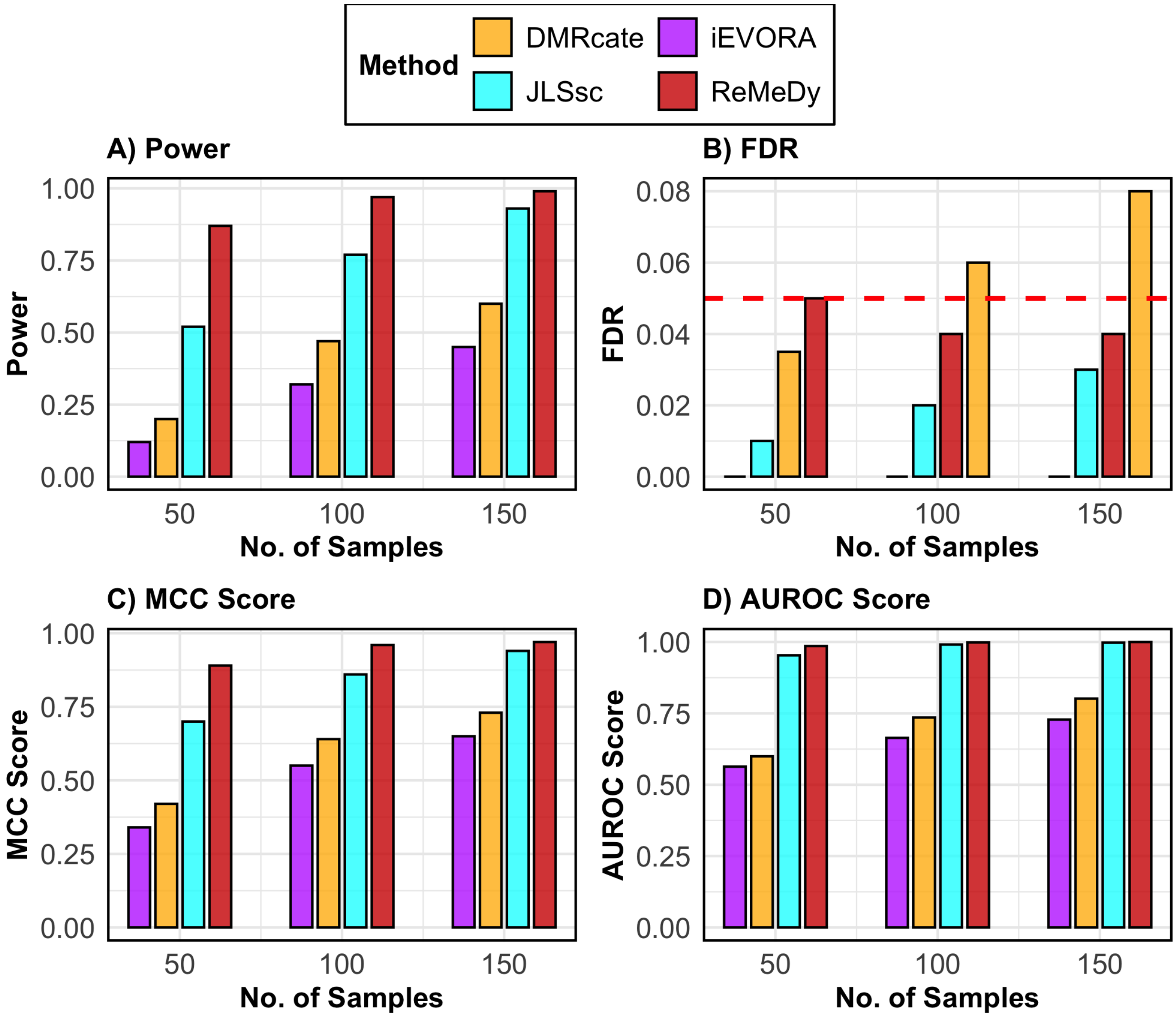

**SFigure** 24: **Performance of ReMeDy and competing methods in the DVMR scenario**. This panel plot compares ReMeDy with other competing methods in DVMR scenario across three sample sizes (50, 100, and 150) with mean effect of 1, variance effect of 2.5 and equal group proportions. Panel A shows statistical power, Panel B shows FDR, Panel C shows the MCC score and Panel D shows AUROC scores. Each bar reflects median of 100 simulation runs. The dashed red line in Panel B marks the nominal FDR threshold of 0.05. Higher values in Panels A, C, and D indicate better performance. Methods without a visible bar achieved value of zero.

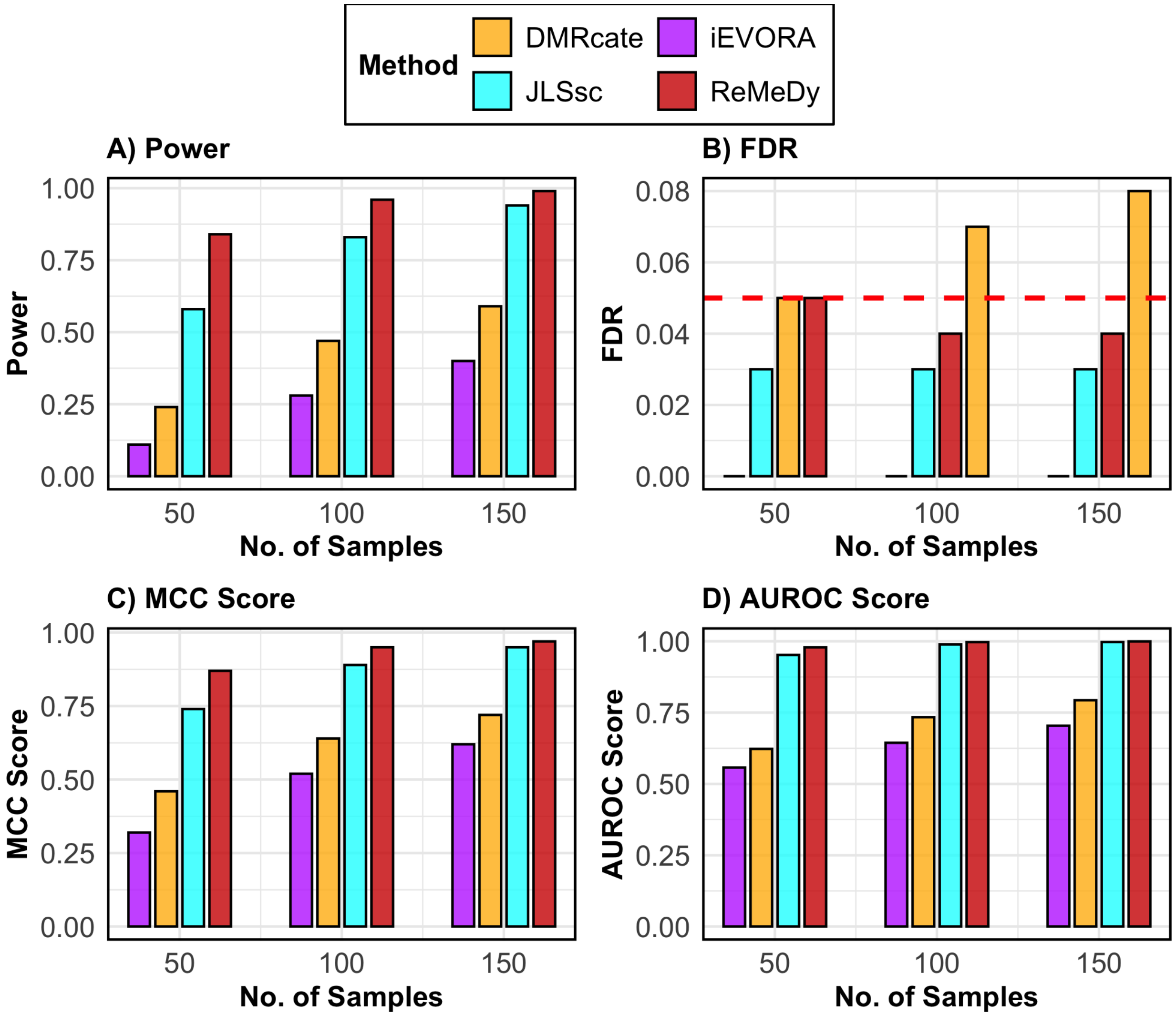

**SFigure 25: Performance of ReMeDy and competing methods in the DVMR scenario**. This panel plot compares ReMeDy with other competing methods in DVMR scenario across three sample sizes (50, 100, and 150) with mean effect of 1, variance effect of 2.5 and unequal group proportions. Panel A shows statistical power, Panel B shows FDR, Panel C shows the MCC score and Panel D shows AUROC scores. Each bar reflects median of 100 simulation runs. The dashed red line in Panel B marks the nominal FDR threshold of 0.05. Higher values in Panels A, C, and D indicate better performance. Methods without a visible bar achieved value of zero.

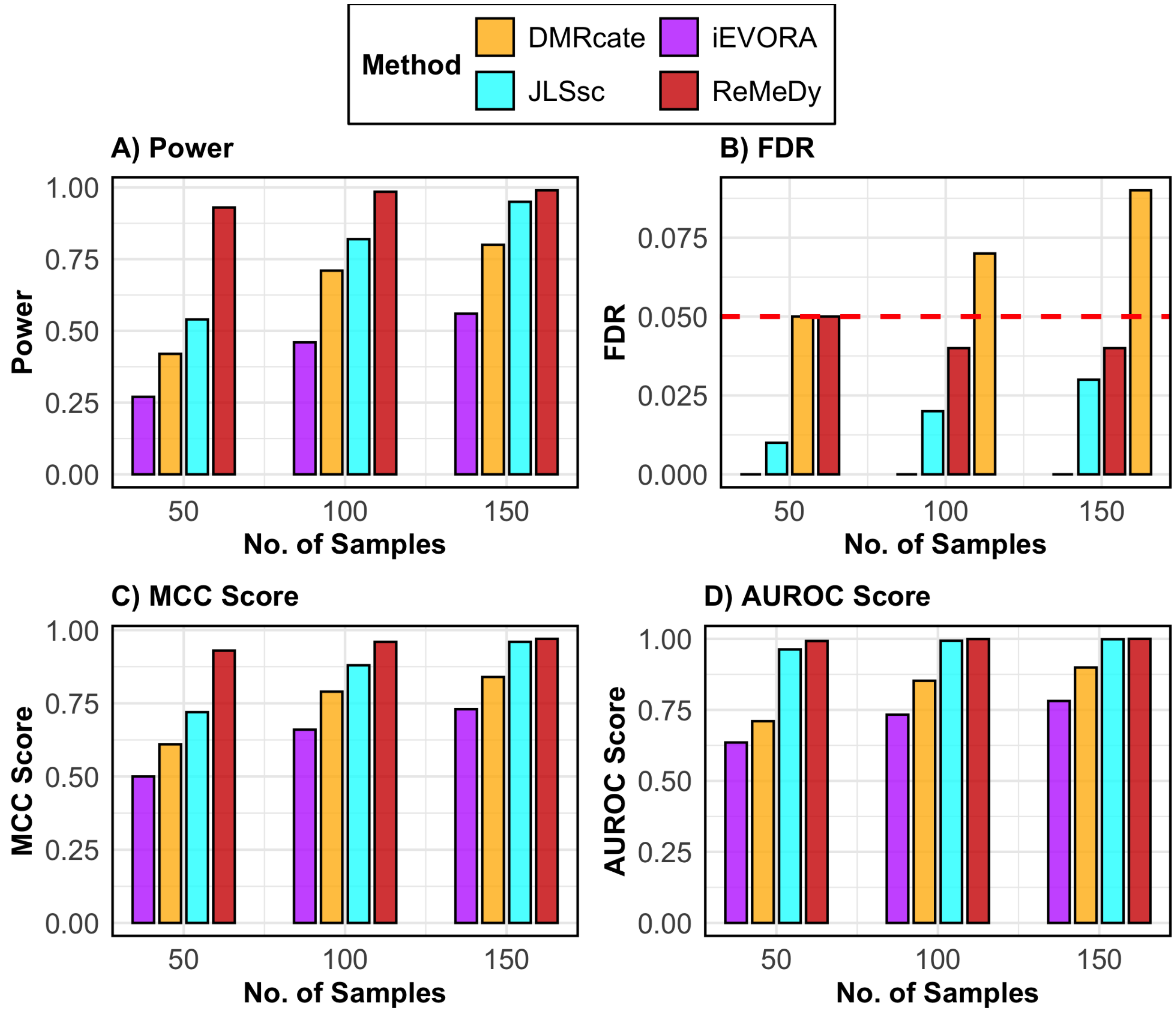

**SFigure 26: Performance of ReMeDy and competing methods in the DVMR scenario.** This panel plot compares ReMeDy with other competing methods in DVMR scenario across three sample sizes (50, 100, and 150) with mean effect of 1, variance effect of 3.5 and equal group proportions. Panel A shows statistical power, Panel B shows FDR, Panel C shows the MCC score and Panel D shows AUROC scores. Each bar reflects median of 100 simulation runs. The dashed red line in Panel B marks the nominal FDR threshold of 0.05. Higher values in Panels A, C, and D indicate better performance. Methods without a visible bar achieved value of zero.

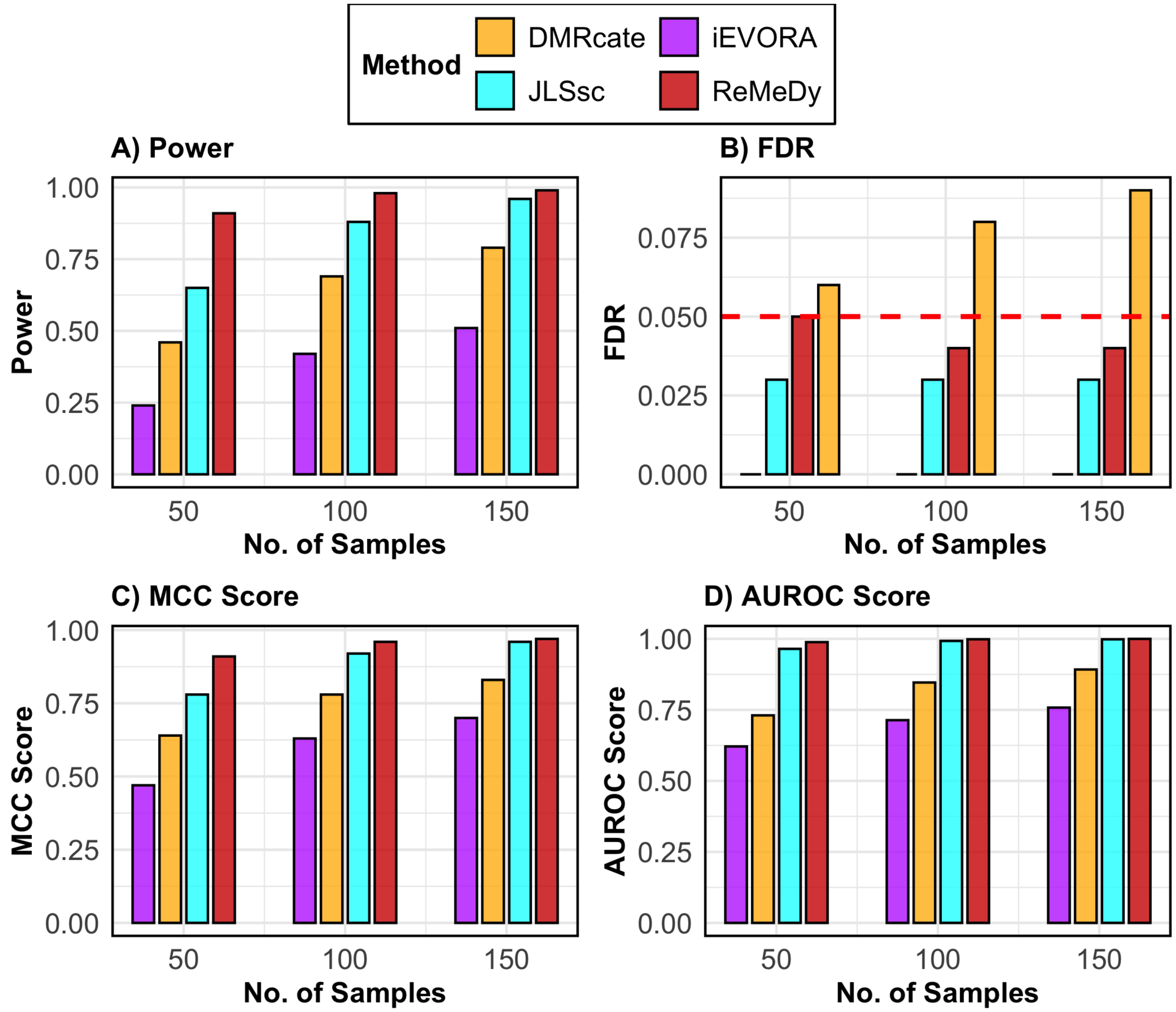

**SFigure** 27: **Performance of ReMeDy and competing methods in the DVMR scenario**. This panel plot compares ReMeDy with other competing methods in DVMR scenario across three sample sizes (50, 100, and 150) with mean effect of 1, variance effect of 3.5 and unequal group proportions. Panel A shows statistical power, Panel B shows FDR, Panel C shows the MCC score and Panel D shows AUROC scores. Each bar reflects median of 100 simulation runs. The dashed red line in Panel B marks the nominal FDR threshold of 0.05. Higher values in Panels A, C, and D indicate better performance. Methods without a visible bar achieved value of zero.

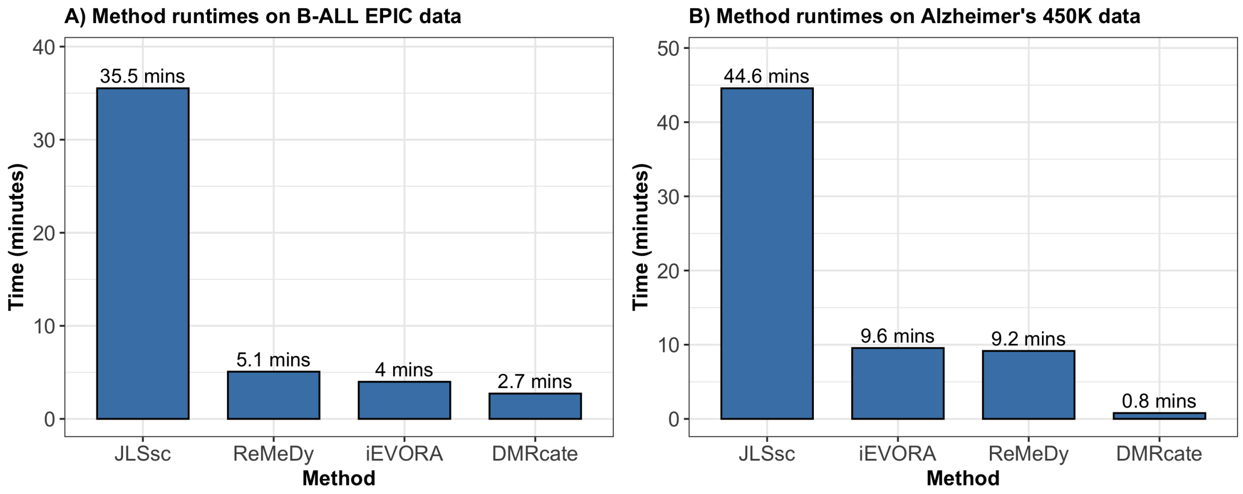

**SFigure** 28: **Runtime comparison of ReMeDy and competing methods on two population-level datasets**. These bar plot compares the computational runtime of ReMeDy with DMRcate, JLSsc and iEVORA on the B-ALL EPIC dataset (Panel A) and Alzheimer's 450K dataset (Panel B). Bars represent the total execution time in minutes for each method and numeric labels above each bar indicate the observed runtime rounded off to 1 decimal. ReMeDy was executed using 8 cores in parallel, whereas the other methods were run using their default settings, as no explicit options for user-defined parallelization were available. Notably the significantly below average time of DMRcate in the Alzheimer's dataset is because it did not find any significant probes, which caused the method to terminate early without executing subsequent steps.

### 3 Supplementary Tables

**STable** 1: **Summary of simulation scenarios evaluated in this study**. This table summarizes the simulation settings used to evaluate method performance under different methylation dysregulation scenarios. The first column lists the scenario types (DMR-only, VMR-only, and DVMR). The second column reports the sample sizes considered. The third and fourth columns specify the mean and variance effect sizes, respectively. The fifth column indicates the group proportions for case–control comparisons. The sixth and seventh columns report the number of regions and CpGs per region used in the simulations. The final column provides the total number of distinct simulation scenarios evaluated for each scenario type.

| Scenario type | Sample sizes | Mean effect | Variance effect | Group proportions (Case-control) | No. of regions | CpGs per region | No. of scenarios |
|---|---|---|---|---|---|---|---|
| DMR-only | 50, 100, 150 | 0.4, 0.7, 1 | 1 | 50/50, 30/70 | 10,000 | 3 | 18 |
| VMR-only | 50, 100, 150 | 0 | 1.5, 2.5, 3.5 | 50/50, 30/70 | 10,000 | 3 | 18 |
| DVMR | 50, 100, 150 | 0.4, 0.7, 1 | 1.5, 2.5, 3.5 | 50/50, 30/70 | 10,000 | 3 | 54 |
| | | | | | | **Total scenarios** | **90** |

[4] Bioconductor Package Maintainer. *TxDb.Hsapiens.UCSC.hg19.knownGene: Annotation package for TxDb object(s)*, 2025. R package version 3.2.2.

[5] Bioconductor Package Maintainer. *TxDb.Hsapiens.UCSC.hg38.knownGene: Annotation package for TxDb object(s)*, 2025. R package version 3.18.0.

[6] John Blair, Ryan Yuen, Brendan Lim, Deborah McFadden, Peter von Dadelszen, and Wendy Robinson. Widespread DNA hypomethylation at gene enhancer regions in placentas associated with early-onset pre-eclampsia. *Molecular Human Reproduction*, 19, 06 2013.

[7] Trevor Breusch and Adrian Pagan. A Simple Test For Heteroscedasticity and Random Coefficient Variation. *Econometrica*, 47, 09 1979.

[8] Morton Brown and Alan Forsythe. Robust Tests for Equality of Variances. *JASA. Journal of the American Statistical Association*, 69, 09 1974.

[9] Rafael G. Cavalcante and M. A. Sartor. *annotatr: Genomic regions in context*, 2017. R package version 1.34.0.

[18] Tamás Lipták. On the combination of independent tests. *Publications of the Mathematical Institute of the Hungarian Academy of Sciences*, 3, 01 1959.

[29] Matthew Suderman, James R Staley, Robert French, Ryan Arathimos, Andrew Simpkin, and Kate Tilling. Dmrff: Identifying differentially methylated regions efficiently with power and control. *bioRxiv*, page 508556, 2018.

[30] Andrew E Teschendorff, Yang Gao, Allison Jones, Matthias Ruebner, Matthias W Beckmann, David L Wachter, Peter A Fasching, and Martin Widschwendter. DNA methylation outliers in normal breast tissue identify field defects that are enriched in cancer. *Nature Communications*, 7(1):10478, 2016.

[31] Andrew E Teschendorff, Francesco Marabita, Matthias Lechner, Thomas Bartlett, Jesper Tegner, David Gomez-Cabrero, and Stephan Beck. A beta-mixture quantile normalization method for correcting probe design bias in Illumina Infinium 450 k DNA methylation data. *Bioinformatics*, 29(2):189–196, 2013.

[32] Paul D. Thomas, Dustin Ebert, Anushya Muruganujan, Tharshana Mushayahama, Laurent P. Albou, and Huaiyu Mi. PANTHER: Making genome-scale phylogenetics accessible to all. *Protein Science*, 31(1):8–22, 2022.

[33] Zongli Xu, Liang Niu, and Jack Taylor. The enmix DNA methylation analysis pipeline for Illumina BeadChip and comparisons with seven other preprocessing pipelines. *Clinical Epigenetics*, 13, 12 2021.

[34] Guangchuang Yu, Li-Gen Wang, Yan Han, and Qing-Yu He. clusterprofiler: an R package for comparing biological themes among gene clusters. *OMICS*, 16(5):284–287, 2012.

[35] Zbyněk Šidák. Rectangular Confidence Region for the Means of Multivariate Normal Distributions. *Journal of The American Statistical Association - J AMER STATIST ASSN*, 62:626–633, 06 1967.
